# Multi-state Continuous-Time Markov Chain Modeling for Chronic Kidney Disease Progression

**DOI:** 10.64898/2026.05.26.727952

**Authors:** Qiumeng Li, Weiqi Chu, Leili Shahriyari

**Author notes:** These authors contributed equally to this work (Weiqi Chu, Leili Shahriyari). Contributing authors.

## Abstract

This paper presents a unified six-state Continuous-Time Markov Chain (CTMC) framework for Chronic Kidney Disease (CKD) progression, with CKD stages 1–5 modeled as transient states and death as an absorbing state. Under a non-homogeneous CTMC formulation, we derive integral representations for transition probabilities, state distributions, sojourn times, and survival-related quantities. We then study the homogeneous case as a tractable baseline and provide explicit formulas for key quantities. Although the methodology is rooted in standard multi-state theory, these expressions are often left implicit in applied analyses; here they are written out explicitly within a unified CKD framework. We construct covariate-dependent transition rates through a proportional hazards structure, using the standard identification of cause-specific hazards with CTMC transition rates. We fit the time-homogeneous baseline model to 335,283 longitudinal observations from 21,100 synthetic electronic health record patients by maximum likelihood. In this synthetic cohort, the covariate model improves held-out log-likelihood per transition over the null model, with stable performance across 10-times-repeated 5-fold cross-validation, and reproduces the main population-level prevalence patterns. The transition-specific estimates can also be translated into sojourn-time and survival summaries. The model suggests that male sex is associated with faster progression across nearly all CKD transitions, and that hypertension shows a stage-dependent association, with lower estimated transition rates in early stages but a substantial acceleration of the Stage 4 to Stage 5 transition. Overall, the proposed framework provides a mathematically explicit approach for studying CKD trajectories from longitudinal health records.

## 1 Introduction

Chronic Kidney Disease (CKD) has emerged as a significant global public health challenge, with a substantial burden of morbidity and mortality (Bikbov et al. 2020). According to the 2024 Clinical Practice Guidelines from Kidney Disease: Improving Global Outcomes (KDIGO), CKD is defined as abnormalities of kidney structure or function that persist for more than three months and affect a person’s health (KDIGO 2024). Diagnosis is based on two main criteria. The first is evidence of kidney damage, such as persistent albuminuria (defined as Urinary Albumin Excretion Rate, AER ≥ 30 mg/24h or Albumin-to-Creatinine Ratio, ACR ≥ 30 mg/g), abnormal urinary sediment, or structural abnormalities identified by imaging. The second is reduced renal function, specifically a Glomerular Filtration Rate (GFR) below 60 mL/min/1.73m^2^ (KDIGO 2024; Levey et al. 2005).

As the central quantitative measure of renal excretory function, GFR represents the total volume of fluid filtered from the glomerular capillaries into Bowman’s capsule per unit time (Stevens and Levey 2009). Although gold-standard measurements of GFR rely on the clearance of exogenous markers such as inulin (*C*_*in*_), these methods are often impractical in large-scale clinical settings because of their complexity and invasiveness. Consequently, estimated GFR (eGFR), derived from endogenous biomarkers such as serum creatinine (SCr) or cystatin C (CysC) through validated empirical equations, has become the standard clinical measure (Webster et al. 2017). The most widely used equations include the MDRD (Modification of Diet in Renal Disease) study equation (Levey et al. 2006) and the CKD-EPI (Chronic Kidney Disease Epidemiology Collaboration) equation (Inker et al. 2021). Based on eGFR values, KDIGO classifies CKD into five stages (*G*1–*G*5), with higher stages corresponding to more severe impairment (KDIGO 2024).

Continuous-Time Markov Chains (CTMCs) provide a natural mathematical framework for modeling disease progression over time (Ross 2014). In medical applications, CTMCs describe how patients move among health states in continuous time. One major advantage of this framework is its ability to accommodate the irregularly spaced follow-up intervals commonly encountered in longitudinal clinical studies and electronic health records (Jackson 2011). CTMC-based models have been applied to problems such as cancer screening (Duffy et al. 1995), diabetes risk prediction (Marshall and Jones 1995), and cognitive decline in Alzheimer’s disease (Zhang et al. 2019). In the context of kidney disease, they offer a useful way to describe the gradual loss of renal function and the stochastic progression toward End-Stage Renal Disease (ESRD) (Bikbov et al. 2020; Eckardt et al. 2013).

Within the CKD literature, several multi-state modeling approaches have been considered. Begun et al. (2013) proposed a non-homogeneous CTMC framework and derived the probability density function for sojourn times under time-dependent transition rates, although their empirical analysis was ultimately simplified to the homogeneous setting. Later, Anwar and Mahmoud (2014) studied a 5-state homogeneous CTMC model and derived analytical formulas for transition probabilities, sojourn times, survival functions, and state probability distributions. These works illustrate both the flexibility of non-homogeneous formulations and the tractability of homogeneous ones, but they also leave room for a more unified exposition linking the two settings within a single CKD progression framework.

This work develops a unified six-state CTMC framework for CKD progression. We first formulate the model in the general non-homogeneous setting, where the law of total probability is used to derive integral representations for transition probabilities, state distributions, sojourn times, and survival-related quantities. We then study the same framework under the time-homogeneous assumption as a tractable baseline. In that setting, the forward Kolmogorov differential equations can be solved directly, yielding explicit expressions for transition probabilities as well as analytical formulas for sojourn times, state probability distributions, survival times, and the expected number of patients in each state. While the underlying methodology is rooted in standard multi-state theory, these quantities are often left implicit in applied analyses; here they are written out explicitly within a unified CKD framework.

To incorporate patient-level covariates, we model each transition rate using a proportional hazards structure. This choice is justified by the standard connection between cause-specific hazards and CTMC transition rates in continuous-time multistate models. As a result, covariates can enter the generator matrix separately for each transition, making their effects transition-specific and interpretable. Although the theoretical framework is presented in both non-homogeneous and homogeneous settings, the numerical analysis in this paper focuses on the time-homogeneous model as a baseline implementation.

We apply the homogeneous baseline model to longitudinal synthetic electronic health record data generated using Synthea v4.0.0 (Walonoski et al. 2018). The model is fitted by Maximum Likelihood Estimation using the msm package (Jackson 2011). We evaluate the fitted model through cross-validation, prevalence calibration, transition-specific parameter estimates, sensitivity analysis, and derived sojourn and survival summaries. Because the analysis is based on synthetic data, the results should be interpreted as a model-based assessment of the proposed framework rather than as clinical evidence about real-world CKD progression. Overall, this paper provides a mathematically explicit CTMC framework for estimating, evaluating, and interpreting CKD progression models from longitudinal health-record data.

## 2 Methods

The clinical condition of patients with CKD typically deteriorates gradually over time. Figure 1 shows the six-state model considered in this paper, in which states 1 to 5 correspond to CKD stages 1 through 5, defined according to the estimated glomerular filtration rate (eGFR). Death is treated as an absorbing state (state 6).

**Fig. 1.**
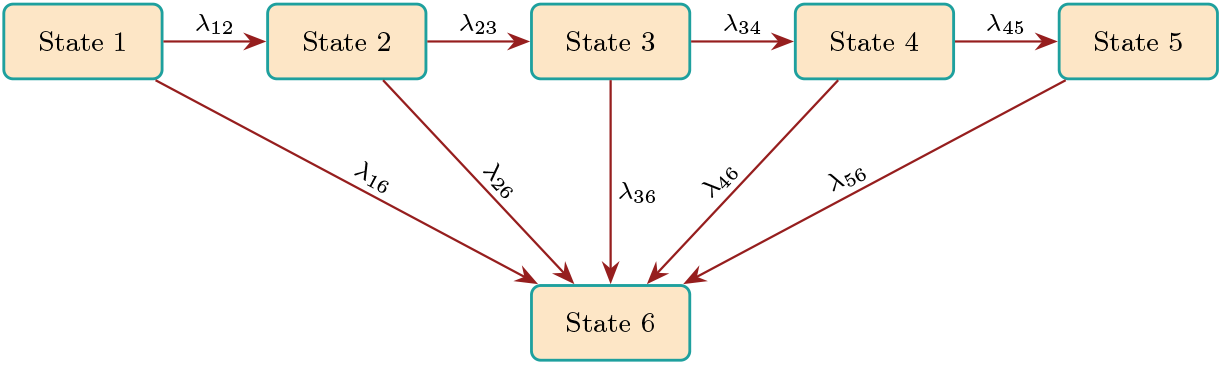
A diagram for the six-state Markov CKD model with transition rates *λ*_*ij*_.

A detailed description of each state is provided in Table 1. We model CKD progression by a non-homogeneous CTMC (*X*_*t*_)_*t*≥0_ defined on the finite state space S = {1, 2, 3, 4, 5, 6}, where *X*_*t*_ denotes the disease state of an individual at age *t*.

**Table 1.** Definitions of states for a six-state CKD model.

| State Number | State Name | eGFR Values (mL/min/1.73m<sup>2</sup>) |
| --- | --- | --- |
| 1 | Normal or high GFR, CKD stage 1 | $\geq 90$ |
| 2 | Mild reduction in GFR, CKD stage 2 | $60 \leq \text{eGFR} < 90$ |
| 3 | Moderate reduction in GFR, CKD stage 3 | $30 \leq \text{eGFR} < 60$ |
| 4 | Severe reduction in GFR, CKD stage 4 | $15 \leq \text{eGFR} < 30$ |
| 5 | Established kidney failure, CKD stage 5 | $< 15$ |
| 6 | Death | – |

### 2.1 Transition rate matrix

Let Λ(*t*) be the age-dependent transition rate matrix (i.e. generator matrix) of the CTMC, where *t* denotes the age. The off-diagonal elements *λ*_*ij*_(*t*) with *i* ≠ *j* are the instantaneous transition rates from state *i* to state *j* at age *t*, defined as

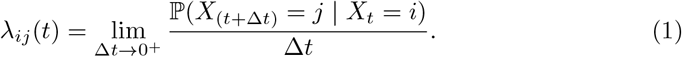

Throughout this paper, we use the term transition rate for *λ*_*ij*_(*t*); this same quantity is also commonly called a transition intensity in the multi-state modeling literature. The total rate of leaving state *i* equals the sum of the transition rates to all other states

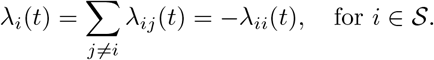

Because CKD is a progressive and generally irreversible disease, we assume that patients transition only to the immediate next disease stage or directly to death, and exclude other transition paths. Therefore, the transition rates satisfy

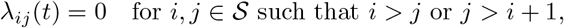

with the exception of transitions to the absorbing death state (*j* = 6). Therefore, the generator matrix Λ(*t*) has the following upper-triangular form

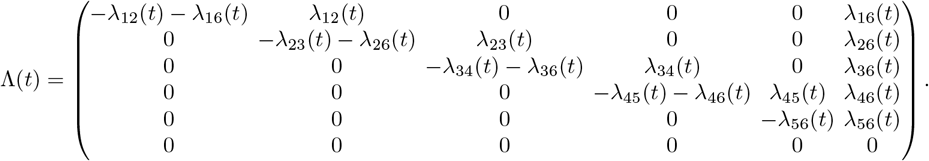

In addition, we define the conditional jump probability *µ*_*ij*_(*t*) as the probability that the process moves to state *j* given that it leaves from state *i*. It is given by

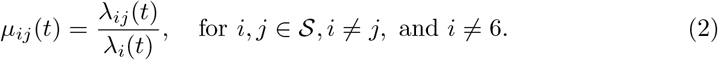

### 2.2 Sojourn time

Let *T*_*i*_(*τ* ) be the holding time that the Markov process stays in state *i*, given that it is in state *i* at age *τ* . That is

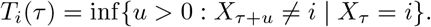

If *τ* is the exact time of entry into state *i*, then *T*_*i*_(*τ* ) is the total holding (sojourn) time in state *i*.

Let *S*_*i*_(*t* | *τ* ) be the state-specific survival function

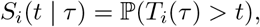

which is the probability that the process remains in state *i* for at least a duration of *t* since the process is in state *i* at age *τ* . Consider an infinitesimally small time increment Δ*t*. By the Markov property, we have

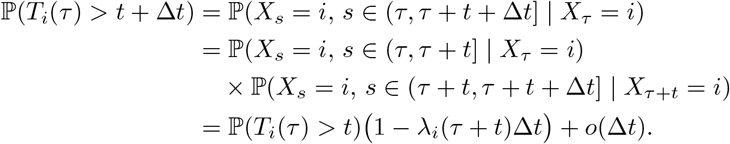

We rearrange terms and divide both sides by Δ*t*, which gives

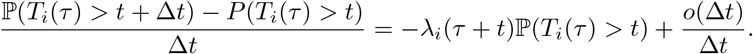

We take the limit Δ*t* → 0^+^ and obtain the first-order differential equation

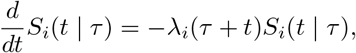

which yields, with the initial condition *S*_*i*_(0 | *τ* ) = ℙ(*T*_*i*_(*τ* ) *>* 0) = 1, that

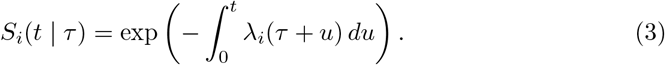

The cumulative distribution function (CDF) of *T*_*i*_(*τ* ) is *F*_*i*_(*t*|*τ* ) = 1 − *S*_*i*_(*t*|*τ*). We take the derivative of the CDF with respect to *t* and obtain the probability density function (PDF) *f*_*i*_(*t* | *τ* ) for the non-homogeneous case

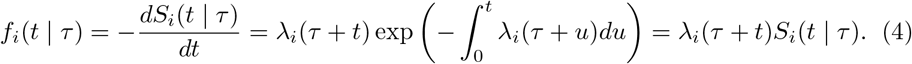

Therefore, if the patient enters state *i* at age *τ*, then the expected sojourn time is

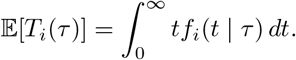

Let *f*_*ij*_(*t* | *τ* ) be the PDF for the event that the process transitions from state *i* to state *j* (where *j* ≠ *i*) after a remaining time *t*, given that it is in state *i* at age *τ* . The PDF *f*_*ij*_ satisfies

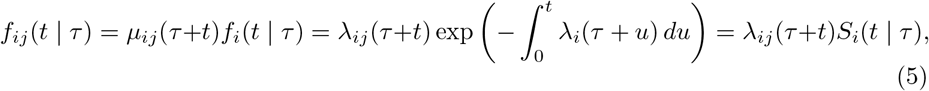

where *µ*_*ij*_ is the probability that the process moves to state *j* given that it leaves from state *i* defined in (2).

In a time-homogeneous CTMC, the transition rates are independent of time (i.e., *λ*_*i*_(*t*) = *λ*_*i*_). We substitute the constant *λ*_*i*_ into Eq. (4) and obtain that the PDF of the holding time *T*_*i*_(*τ* ) is an exponential distribution, which is

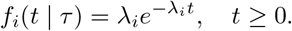

Consequently, the expected sojourn time in state *i* is simply the reciprocal of the total transition rate out of that state, which is

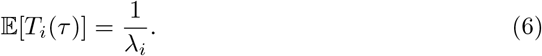

For our six-state CKD model, the expected sojourn times are

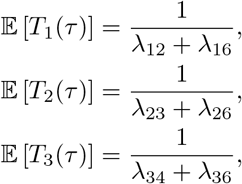

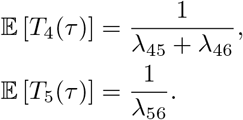

### 2.3 Forward Kolmogorov differential equations

Let *P*_*ij*_(*τ, t*) be the transition probability function, which is the probability that the process is in state *j* at time *t*, given that it was in state *i* at time *τ* . From the six-state transition paths (i.e. no backward transitions), we can represent *P*_*ij*_ in the upper-triangular matrix form, which is

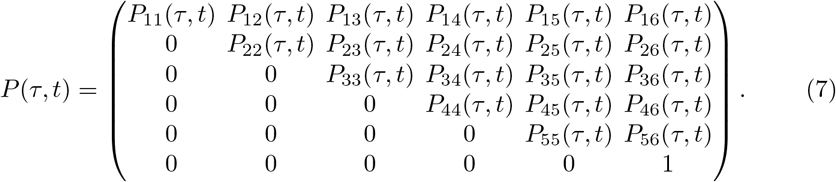

Due to the definition of transition probabilities, each row sum equals 1 for any *t* and 0 ≤ *τ* ≤ *t*, and the initial conditions at *t* = *τ* are

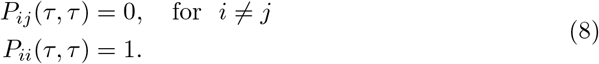

Following the standard derivation of the Kolmogorov forward equations (Norris 1997), we apply the Chapman–Kolmogorov identity to the infinitesimal transition probabilities and take the limit Δ*t* → 0^+^. This gives

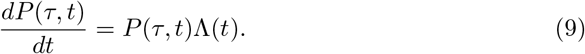

Together with the initial condition *P* (*τ, τ* ) = *I*, where *I* is the 6 × 6 identity matrix, this system determines the transition probability matrix *P* (*τ, t*).

### 2.4 Transition probability functions

For the non-homogeneous CTMC, if a time-dependent generator Λ(*t*) commutes with its integral 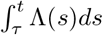, the Kolmogorov forward equation (9) yields an explicit solution

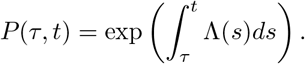

In CKD progression models, however, age-dependent or covariate-dependent generators need not satisfy this commutativity condition. In this case, we should solve the Kolmogorov forward equations numerically. A straightforward computational approach is numerical integration of the matrix differential equation, for example, by Runge–Kutta methods. Since the state space in our CKD model is small, this approach is easy to implement and well-suited for obtaining transition probabilities numerically. We also develop a path-based integration formulation. Its main advantage is not computational simplicity but interpretability: it decomposes transition probabilities into contributions from possible disease-progression paths, makes the role of intermediate stages explicit, and provides a natural foundation for likelihood contributions under partially observed trajectories and for related quantities such as sojourn times and survival probabilities.

Here, we introduce the path-based integration formulation from the standard sample-path construction of CTMCs (Norris 1997). Since the disease progression transitions only to the immediate next state or death, transitioning from state *i* to state *j*≠ 6 follows a single valid chronological pathway: *i* → *i* + 1 → · · · → *j*. Let the continuous random variable *s*_*k*_ (*k* ∈ {*i* + 1, …, *j*}) denote the exact age of entry into state *k*, with initial age *s*_*i*_ = *τ* . All intermediate transition times form an ordered time simplex Ω_*ij*_,

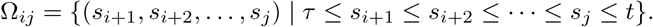

By the Markov property, the joint density of this unique transition pathway factorizes as

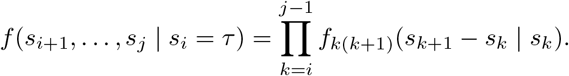

For any given state *i*, the probability of staying in state *i* during the time interval (*τ, t*], given by the survival function (3), is

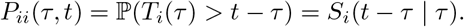

Therefore, the probability that the process enters state *j* at age *s*_*j*_ and remains in state *j* until age *t* is *S*_*j*_(*t* − *s*_*j*_ |*s*_*j*_).

We apply the continuous law of total probability and multiply the joint probability density of the pathway by the final survival probability. We integrate over the entire ordered time simplex Ω_*ij*_ and obtain

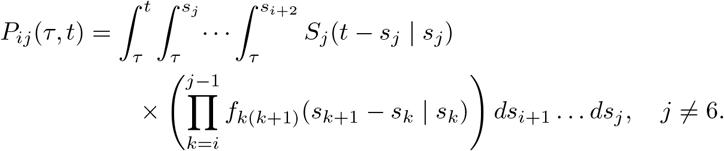

Since state 6 is an absorbing state, its survival probability is always 1, i.e. *S*_6_(*t* − *s*_6_ | *s*_6_) = 1. According to our assumption, an individual directly faces a fatal risk *λ*_*j*6_(*t*) and transitions into state 6 from any prior transient state *j* (*j* ∈ {*i, i* + 1, …, 5}). Therefore, reaching state 6 from state *i* is the union of multiple mutually exclusive jump pathways. Let the final jump to the absorbing state occur from state *j*; this specific pathway is *i* → *i* + 1 → · · · → *j* → 6. Under this specific pathway, the individual undergoes a sequential progression from *i* to *j*, and subsequently enters the absorbing state at time *s*_6_. Incorporating the survival probability *S*_6_(*t* ∈ *s*_6_ *s*_6_) = 1, the probability integral for this specific trajectory is

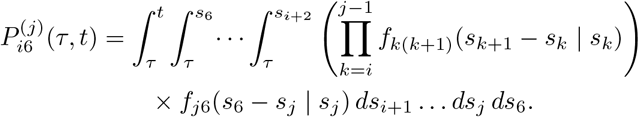

Because the fatal events originating from different transient states *j* are mutually exclusive, the law of total probability dictates that the total probability of reaching the absorbing state 6 by time *t* is the sum of the probabilities of all possible pre-jump states *j*, which is

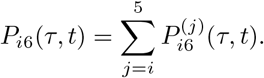

Substituting the integral form of 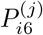 yields the final analytical expression for the generalized absorbing state probability

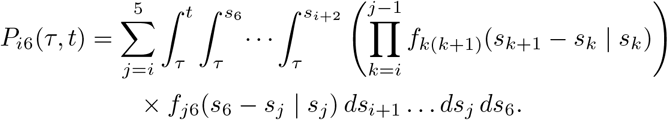

Under the time-homogeneous assumption, the transition rates are constant over time, and the solution of the transition probability function *P* (*τ, t*) admits the Taylor series expansion

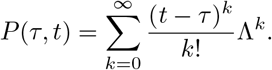

In addition, one can derive an analytical expression for each element of *P* (*τ, t*) by directly solving the forward Kolmogorov differential equations (9). When the exit rates *λ*_*i*_, …, *λ*_*j*_ are distinct, we have the solution

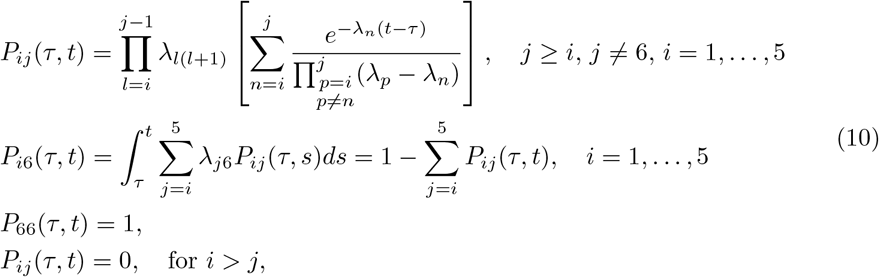

where *λ*_*i*_ = ∑_*j*≠*i*_ *λ*_*ij*_ for *i* 1,, 5 . When two exit rates coincide, *λ*_*i*_ = *λ*_*j*_, the solutions to the forward Kolmogorov equations (9) are obtained by taking the limit *λ*_*i*_ → *λ*_*j*_ in the general formula (10). More generally, when multiple exit rates are equal, the solutions remain analytically valid, but the explicit formulas depend on the multiplicity pattern of the repeated rates and must be handled on a case-by-case basis. In all cases, the system (9) can be solved either analytically or numerically. In Appendix A, we spell out the formulas for *P*_*ij*_ in (10) and provide an illustrative example with equal exit rates.

### 2.5 State probability distribution

To analyze the progression of the CKD process, it is essential to estimate the state probability distribution at a specific time *t*, denoted by the row vector **p**(*t*), where the *i*th entry *p*_*i*_(*t*) is the probability that an individual is in state *i* at age *t*. The evolution of this distribution is governed by

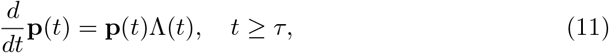

with the initial condition at starting age *τ* given by **p**(*τ* ) with *p*_6_(*τ* ) = 0, assuming the individual has not entered the absorbing death state at the initial time.

For any state *j*, by the law of total probability, we have

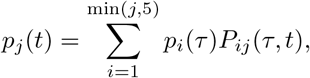

where *P*_*ij*_ is the transition probability defined in (7). The probability *p*_*j*_(*t*) has two sources of contributions: the probability that an individual initially in state *j* remains there until time *t*, and the sum of probabilities of entering state *j* from any preceding state *i < j*. We express this relation as

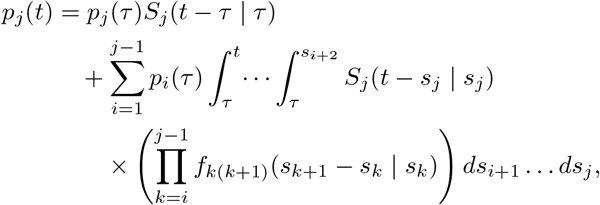

where *f*_*ij*_(*t*|*τ* ) = *λ*_*ij*_(*τ* + *t*)*S*_*i*_(*t*|*τ* ) is defined in Eq. (5) For the absorbing state (*j* = 6), we have

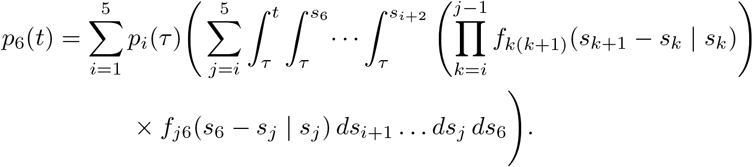

Under the time-homogeneous CTMC assumption, where transition rates are constant, solving the Kolmogorov forward differential equations (11) yields explicit analytical expressions for the state probabilities. If the exit rates *λ*_*i*_, …, *λ*_*j*_ are distinct, the transient-state probability is

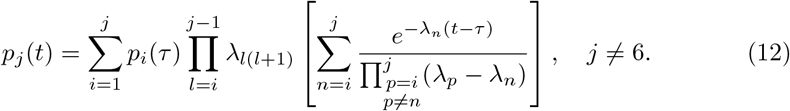

If some of these exit rates are equal, the transition probabilities remain well defined, but the compact formula above is no longer valid as written. The corresponding repeated-rate expressions can be obtained either by solving the forward equations (11) directly or by taking the appropriate limits of the distinct-rate formula (12). Further details are provided in Appendix B.

### 2.6 Model-based expected time to absorption

For an individual in the CKD progression model, we define *U*_*i*_(*τ* ) as the time until absorption (death), given that the individual enters state *i* at age *τ*, where *i* ∈ {1, …, 5}. Let *T*_*i*_(*τ* ) be the sojourn time in state *i*, and let *J*_*i*_ denote the destination state of the next transition after leaving state *i*. Because the model allows either progression to the next CKD stage or direct transition to death, the time to absorption satisfies

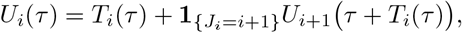

where the continuation term is zero if the next transition is directly to death. Taking expectations and conditioning on the duration *T*_*i*_(*τ* ) = *t* and the next destination, we obtain

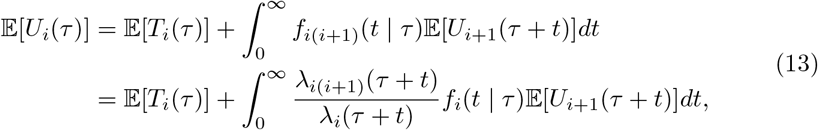

using the relation of the transition density *f*_*ij*_(*t*|*τ* ) in Eq. (5).

Under the time-homogeneous assumption, we have 𝔼[*U*_*j*_(*t*)], *λ*_*i*_(*t*), and *λ*_*ij*_(*t*) all independent of age *t*, and 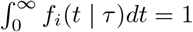. Using 𝔼[*T*_*i*_] = 1*/λ*_*i*_ from Eq. (6), we can further simplify (13) as

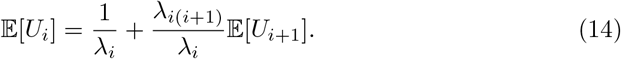

With 𝔼[*U*_6_] = 0, we can calculate the expected time to absorption from each state *i* ∈ {1, …, 5} from (14) in the reverse order, which is

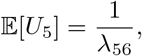

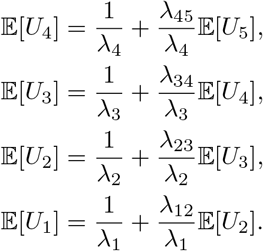

### 2.7 Expected number of patients in each state

The expected number of patients in each state at age *t* can provide useful model-based summaries for resource planning in real-data applications. Let **M**(*t*) ∈ ℝ^6^ be a row vector, where *M*_*j*_(*t*) denotes the number of patients in state *j* at age *t*. Let K be the set of all patients in the system. For each patient *k* ∈ 𝒦, let *τ*_*k*_ denote the age at the most recent observation prior to *t*, and let *s*_*k*_ ∈ {1, …, 6} denote the observed state at that time. We define the indicator variable 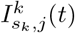 by

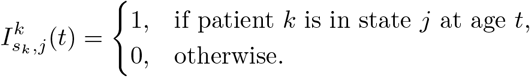

The expectation of this indicator equals the corresponding transition probability

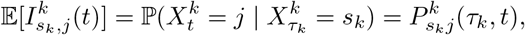

where 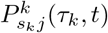 is the probability that patient *k* transitions from state *s*_*k*_ at age *τ*_*k*_ to state *j* at age *t*, which we define in Eq. (7). By the linearity of expectation, we have

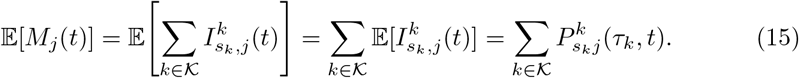

We note that the transition probabilities 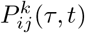 may vary across patients through individual-specific characteristics, such as sex or baseline clinical conditions, reflected in non-homogeneous transition rates.

### 2.8 Equivalence between hazard functions and transition rates

The transition rates *λ*_*ij*_(*t*) in (1) can be related to the cause-specific hazard from survival analysis. Let *T* denote the random time of transition out of the current state *i*, and let *D* denote the destination state. The cause-specific hazard for a transition from state *i* to state *j* is defined as

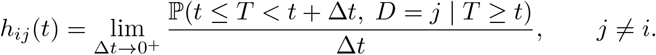

This quantity has the same infinitesimal interpretation as the transition rate *λ*_*ij*_(*t*) defined in (1): both are defined through the conditional probability, per unit time, of moving from state *i* to state *j*. Therefore, under the standard continuous-time multi-state formulation,

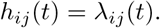

This identification provides the link between survival-analysis notation and the generator-based CTMC formulation, and motivates the use of transition-specific proportional-hazards structures within the generator matrix (Andersen et al. 1993, 2002).

Building upon this foundation, we explicitly parameterize the transition rates to model the dynamics of disease progression. Following Gompertz’s observation that adult mortality increases exponentially with age (Gompertz 1825), we extend this principle to our CTMC framework. We assume that the transition rate *λ*_*ij*_(*t*; *z*) from state *i* to state *j* depends exponentially on the subject’s age *t* and an observed column vector of covariates *z*, which yields

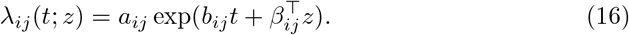

Here, *a*_*ij*_ *>* 0 is the baseline scale parameter, and *b*_*ij*_ is a scalar capturing the age dependency. The term *β*_*ij*_ represents a vector of regression coefficients corresponding to the explanatory factors in *z* (e.g., sex, hypertension status).

This parametric form translates physiological aging into mathematical constraints. For a positive *b*_*ij*_, the model reflects a decline in vitality with age (Yashin et al. 2007). While the Gompertz form is traditionally used to model overall mortality, the same exponential-in-age structure can be used as a flexible parametric specification for stage-specific transition rates. Furthermore, setting *b*_*ij*_ = 0 accommodates a constant baseline rate as a special case, reducing the model to a time-homogeneous Markov process.

The dependency on covariates *z* is incorporated via a proportional hazards structure (Cox 1972), ensuring the interpretability of epidemiological risk factors. While (16) focuses on observed heterogeneity, this framework can be extended with frailty terms to account for unobserved heterogeneity (Vaupel et al. 1979).

### 2.9 Maximum likelihood estimation

Let *ω* ∈ Ω denote the unknown parameter vector that parametrizes the transition rates *λ*_*ij*_(*t*), including baseline scales, age-dependent components, and covariate effects. To estimate *ω* for the proposed multi-state model, we employ the Maximum Likelihood Estimation (MLE) framework.

In practice, the likelihood function depends on the observation scheme of the longitudinal data. To make this dependence explicit, we distinguish three common data structures encountered in clinical studies. Assuming independence across patients, the likelihood can be written as a product of contributions from the observed data:

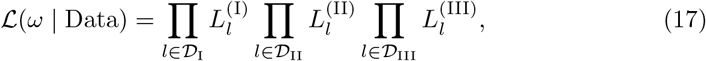

where 𝒟_I_, 𝒟_II_, and 𝒟_III_ denote the sets of observations under the three data structures, and *L*^(I)^, *L*^(II)^, and *L*^(III)^ are the corresponding likelihood contributions.

The first type of data set (𝒟_I_) assumes full observation. For each data point collected, we know the exact age *τ* when the patient enters state *i* and their precise sojourn time *t* before transitioning to state *j*. This single transition contributes to the likelihood function in the form of the cause-specific probability density function *f*_*ij*_(*t* | *τ* ) in (5), which is

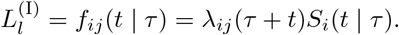

The second type of data set is called the panel data (𝒟_II_), which assumes partial observations at each individual level. In this scenario, we collect individual states at discrete ages, while lacking information regarding the exact intermediate states or the precise arrival and departure times of states. Suppose we observe that a patient is in state *i* at age *τ* and in state *j* at age *t*. Then, this data point contributes to the likelihood function in the form of the transition probability *P*_*ij*_(*τ, t*) (Eq. (7)), which is

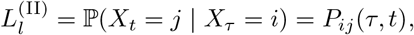

where *P*_*ij*_(*τ, t*) depends on the parameters (e.g. *λ*_*ij*_(*t*)) and can be solved from the Kolmogorov forward equations (9) as detailed in Section 2.4.

The third type of data set (𝒟_III_) contains the exact arrival times. In this case, we know the corresponding state *i* at age *τ*, and the exact transition time *t* to the destination state *j*. However, the intermediate states remain hidden. The likelihood contribution represents the marginal density of arriving exactly at time *t*. Setting the final arrival time *s*_*j*_ = *t*, we integrate the joint path density over all possible intermediate transition ages (*s*_*i*+1_, …, *s*_*j*−1_)

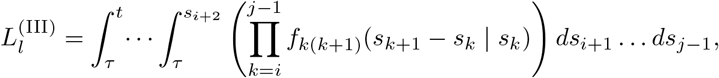

where the initial age is *s*_*i*_ = *τ*, and an empty product evaluates to 1.

In this study, our clinical observations are recorded at discrete, irregular intervals without exact transition times. This cohort configuration corresponds exclusively to the second type, panel data. Accordingly, the likelihood used in our empirical analysis is built entirely from the transition probabilities *P*_*ij*_(*τ, t*) derived from the generator matrix Λ(*t*).

## 3 Application to a synthetic CKD cohort

We apply the proposed CTMC framework to synthetic electronic health record (EHR) data generated by Synthea v4.0.0 (Walonoski et al. 2018), an open-source patient population simulator that produces realistic longitudinal clinical records. From a simulated population of 229,989 patients, 21,100 (9.2%) receive a CKD diagnosis and constitute the analytic cohort. The eGFR thresholds in Table 1 are used only to map each observation within this already-CKD cohort to one of the six CTMC states; CKD diagnosis itself is taken directly from the Synthea diagnosis record rather than being inferred from eGFR alone.

The longitudinal records are processed into a panel suitable for multi-state CTMC analysis through eGFR-to-stage mapping, monotonicity enforcement, six-month thinning with transition preservation, and the addition of death rows for the 7,227 deceased patients (34.3%); full details are given in Appendix C. The final analytic dataset comprises 335,283 person-observations from 21,100 patients.

For the illustrative model considered here, we fix time-invariant covariates at the values recorded at each patient’s first clinical encounter. These include sex (binary indicator: 1 = male, 0 = female) and hypertension (binary indicator: 1 = diagnosed at or before the first clinical encounter, 0 otherwise), which are used in the covariate vector *z* in (16). Synthea generates all CKD patients through a metabolic syndrome or diabetic pathway, so diabetes is present in 100% of the CKD cohort and provides no discriminating power; we therefore exclude it from the covariate vector *z* in (16). We also exclude urine protein concentration from the covariate vector because it is mechanistically determined by the CKD stage within the Synthea model, thus offering no independent clinical signal beyond the state variable itself. The characteristics of the final analytic cohort are summarized in Table 2.

**Table 2.** Characteristics of the analytic cohort (*n* = 21,100).

| Characteristic | Value |
| --- | --- |
| Male, $n$ (%) | 12,668 (60.0%) |
| Female, $n$ (%) | 8,432 (40.0%) |
| Hypertension, $n$ (%) | 18,355 (87.0%) |
| — Male | 11,169 (60.8% of HTN) |
| — Female | 7,186 (39.2% of HTN) |
| Deceased during follow-up, $n$ (%) | 7,227 (34.3%) |
| Total person-observations | 335,283 |

## 4 Numerical illustration under the homogeneous model

Using the msm package (Jackson 2011), we fit the age-independent baseline transition rates *a*_*ij*_ and the covariate regression coefficients *β*_*ij*_ in (16) (with *b*_*ij*_ = 0 under the homogeneous assumption) for the binary covariates of sex and hypertension, as specified in the homogeneous CTMC model above. Because both covariates are fixed per patient, the transition rate *λ*_*ij*_(*z*_*i*_) remains constant over time for each individual, so the fitted process is strictly time-homogeneous with Λ(*t*) ≡ Λ. The remainder of this section reports five aspects of the fitted model: overall fit, prevalence calibration, transition-specific parameter estimates, sensitivity of the least stable coefficient, and clinically interpretable quantities such as sojourn times and survival outcomes.

### 4.1 Assessing overall model fit

As a practical check on model adequacy, we perform 10-times-repeated 5-fold cross-validation (CV), yielding 50 train/test splits in total, to quantify the predictive gain from the covariates and assess the model’s generalization performance. We use stratified splits based on the initial CKD stage to ensure each fold is representative; for each of the 10 repeats a fresh random partition is drawn. In each fold, we fit the full model (*M* 1) and a null model (*M* 0) with 80% of the patients (training set) and evaluate on the held-out 20% (test set). *M* 1 is the full model with sex and hypertension as covariates, i.e., *β*_*ij*_ freely estimated for all transitions (*i, j*) in (16); *M* 0 is the null model without covariates, i.e., *β*_*ij*_ = 0 for all (*i, j*).

The primary metric is the log-likelihood per transition, denoted *LL/n*, where *LL* = log ℒ (*ω* |Data) is the log-likelihood in (17) evaluated on the panel data, and *n* is the total number of observed consecutive state pairs. Higher values of *LL/n* indicate better predictive fit.

As shown in Figure 2, the full model *M* 1 consistently achieves higher held-out *LL/n* values than the null model *M* 0 across the 50 splits (*M* 1 exceeds *M* 0 in all 50 splits). The mean held-out *LL/n* is −0.5641 for *M* 1 and −0.5686 for *M* 0, with a split-to-split standard deviation of 0.0048 for both models. A one-sided paired Wilcoxon signed-rank test (Demšar 2006) on the 50 held-out *LL/n* pairs supports the stability of this advantage across the repeated splits (*p* ≈3.9 × 10^−10^). We note that the 50 splits in repeated cross-validation are not fully independent, since patients reappear across test folds; this *p*-value should therefore be interpreted as supportive evidence of a stable improvement rather than as a strict independent-sample significance test. Although the improvement is modest in absolute magnitude, it is stable across the repeated partitions, suggesting that sex and hypertension provide additional predictive information beyond the baseline transition rates.

**Fig. 2.**
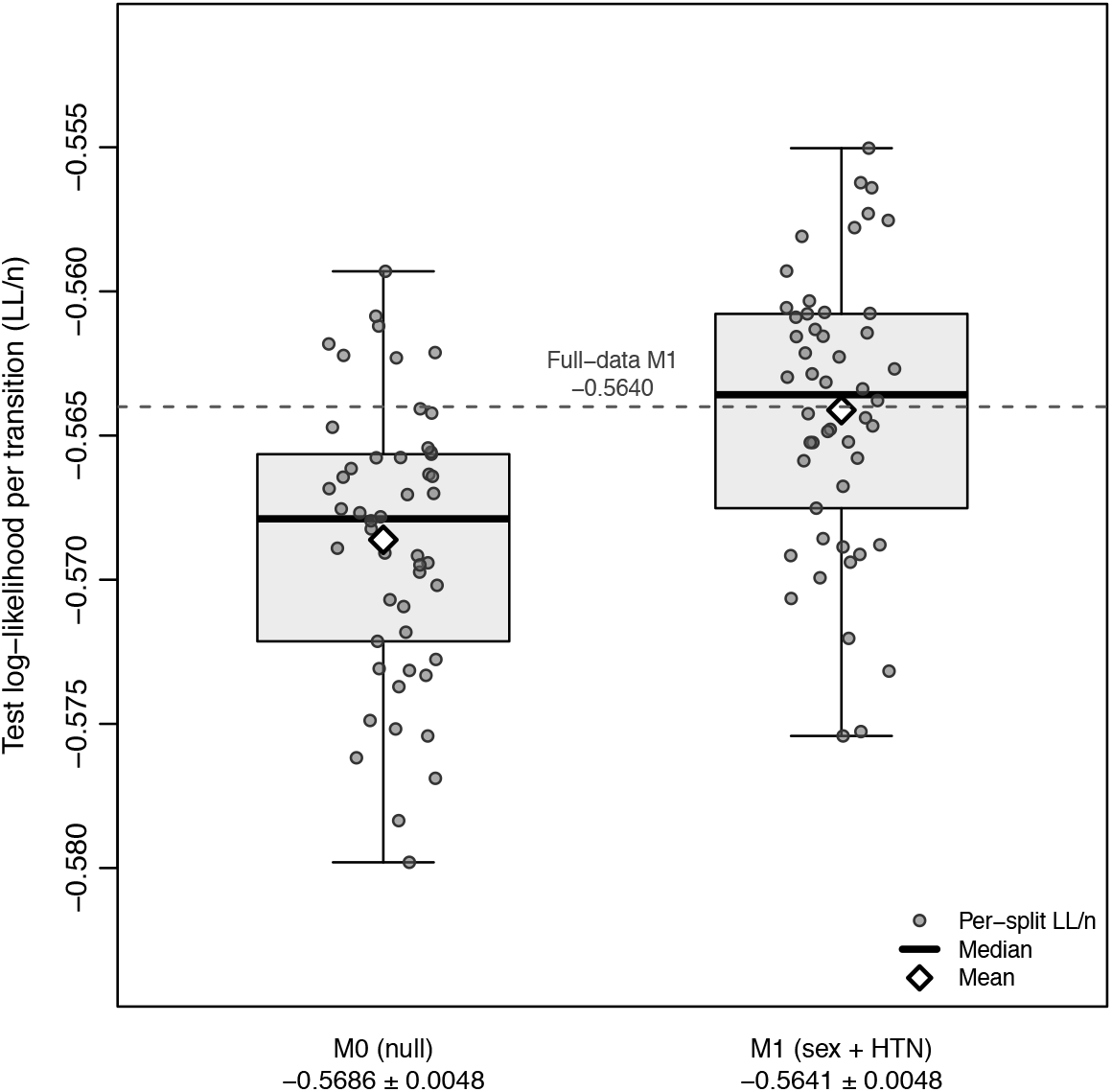
Log-likelihood per transition (*LL/n*) from repeated 5-fold cross-validation for *M* 0 and *M* 1. Jittered dots show individual held-out splits, open diamonds mark the means, and the dashed grey line marks the full-data *M* 1 value (*LL/n* = −0.5640).

The held-out performance of *M* 1 is also very close to its full-data value (*LL/n* = − 0.5640), indicating that the fitted model generalizes well to unseen patient transitions. Together with the large sample-to-parameter ratio (over 3 × 10^5^ observed transitions for 27 free parameters), this provides little evidence of substantial overfitting.

### 4.2 Population-level prevalence and model validation

We next examine how the fitted model predicts population-level state prevalences between ages 40 and 85. For each target age *t* ∈ {40, 45, …, 85}, we define the evaluable subset 𝒦_*t*_ ⊆ 𝒦 of patients who have a usable baseline observation prior to age *t* and have not been lost to follow-up. We use the expected patient counts 𝔼[*M*_*j*_(*t*)] derived in Equation (15) and calculate the model-predicted prevalence of state *j* as 𝔼 [*M*_*j*_(*t*)]*/* 𝒦_*t*_ . To assess population-level calibration, we compare these model-predicted prevalences with the empirical prevalences computed directly from the observed data. The empirical prevalence of state *j* at age *t* is the proportion of 𝒦_*t*_ in state *j*, and the model-predicted prevalence averages each patient’s predicted transition probability into state *j* over the same 𝒦_*t*_. The shared denominator |𝒦_*t*_| makes the two directly comparable. The detailed construction of 𝒦_*t*_ (window width, lost-to-follow-up criterion, LOCF rule, handling of deaths) is given in Appendix C. Figure 3 compares the empirical state prevalences with the model-predicted prevalences from both the model fitted on 100% of the data and the out-of-sample predictions averaged across the 50 splits of the repeated 5-fold cross-validation.

**Fig. 3.**
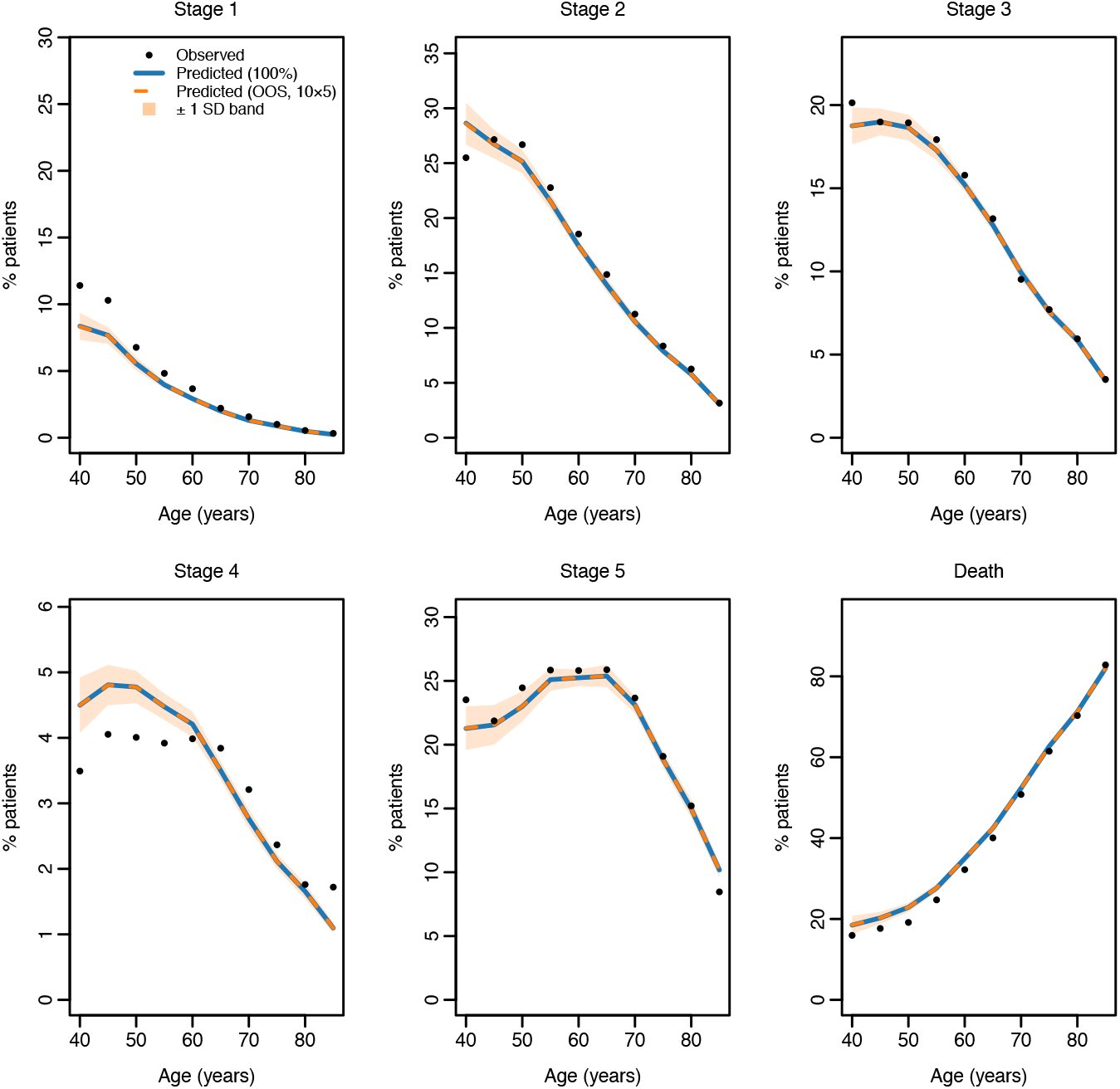
State prevalences at 5-year intervals from age 40 to 85. Observed (solid circles): empirically observed state proportions from the data. Predicted (100%, solid line): expected prevalences from the model fitted on 100% of the data. Predicted (OOS, dashed line): out-of-sample expected prevalences from 10-times-repeated 5-fold cross-validation, where each split trains on 80% of patients and predicts state occupancy for the held-out 20%; the dashed line shows the mean across the 50 splits and the shaded band is ±1 SD across splits.

Overall, the fitted homogeneous CTMC captures the main age-related patterns in the empirical prevalence curves. The predicted prevalence based on the full data set is very close to the OOS prediction averaged across the 50 splits, indicating that the prevalence estimates are relatively stable. For Stages 2, 3, 5, and Death, the predicted curves closely track the observed proportions across most ages, suggesting that the homogeneous CTMC captures the main population-level patterns of disease progression and mortality in this cohort.

However, some deviations are still visible, especially at earlier ages. For example, the model slightly underpredicts Stage 1 prevalence near age 40 and does not fully match the observed Stage 4 pattern. This may be partly due to the time-homogeneous assumption. Because the model uses the same transition rates for all ages, it tends to capture the average progression pattern over the whole age range, but may miss some age-specific changes in the observed prevalence curves.

### 4.3 Parameter estimates and hazard ratios

We now turn to the fitted transition-specific parameters. Table 3 reports the estimated baseline rates *a*_*ij*_ with 95% confidence intervals. Under the proportional hazards specification, these baseline rates describe the transition rates for the reference covariate profile (female, no hypertension). The estimates confirm that disease progression primarily follows the sequential Stage 1 → 2 → 3 → 4 → 5 → 6 sequence, while direct mortality from earlier transient states remains comparatively rare.

**Table 3.** Estimated baseline transition rates *a*_*ij*_ with 95% confidence intervals.

| transition | $\hat{a}_{ij}$ | CI lower | CI upper |
| --- | --- | --- | --- |
| 1 → 2 | 0.5423 | 0.5036 | 0.5839 |
| 1 → 6 | 0.0014 | 0.0004 | 0.0054 |
| 2 → 3 | 0.1665 | 0.1567 | 0.1770 |
| 2 → 6 | 0.0149 | 0.0121 | 0.0183 |
| 3 → 4 | 0.1927 | 0.1810 | 0.2052 |
| 3 → 6 | 0.0207 | 0.0170 | 0.0252 |
| 4 → 5 | 0.2885 | 0.2707 | 0.3076 |
| 4 → 6 | 0.0166 | 0.0126 | 0.0218 |
| 5 → 6 | 0.0746 | 0.0680 | 0.0818 |

Table 4 reports the estimated hazard ratios 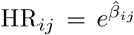 for each covariate and transition, together with 95% confidence intervals. Since both covariates are binary (sex: 1 = male, 0 = female; hypertension: 1 = diagnosed, 0 = no diagnosis), each hazard ratio HR_*ij*_ represents the multiplicative change in transition rate for the group coded as 1 relative to the reference group (female or no hypertension, respectively). A value greater than 1 indicates a higher transition rate for males (or hypertensive patients), and a value below 1 indicates a lower rate relative to the reference group. Bold entries have 95% confidence intervals for the hazard ratios that exclude 1, indicating that the coefficients *β*_*ij*_ are significant, with 0 not in their 95% confidence intervals. To visualize these effects and assess their stability, Figures 4a and 4b display the hazard ratios as forest plots. These plots overlay the full-model point estimates (black filled circles) with the cross-validation split-level (50 splits from 10 × 5-fold CV) means ± 1 SD (open blue diamonds and whiskers).

**Table 4.**
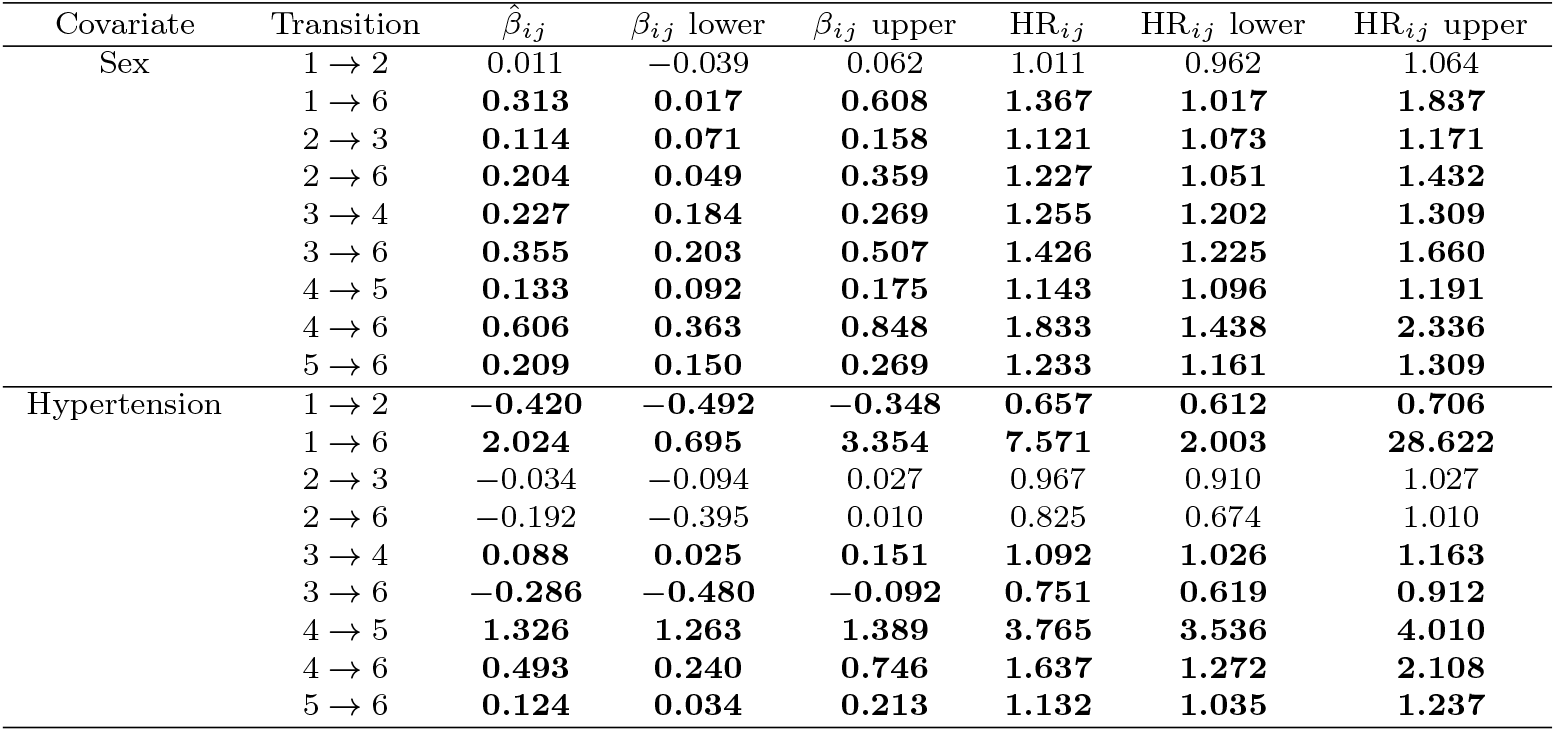
Estimated coefficients *β*_*ij*_ and hazard ratios HR_*ij*_ for each covariate and transition. Bold entries indicate 95% CI of the hazard ratios exclude 1.

**Fig. 4.**
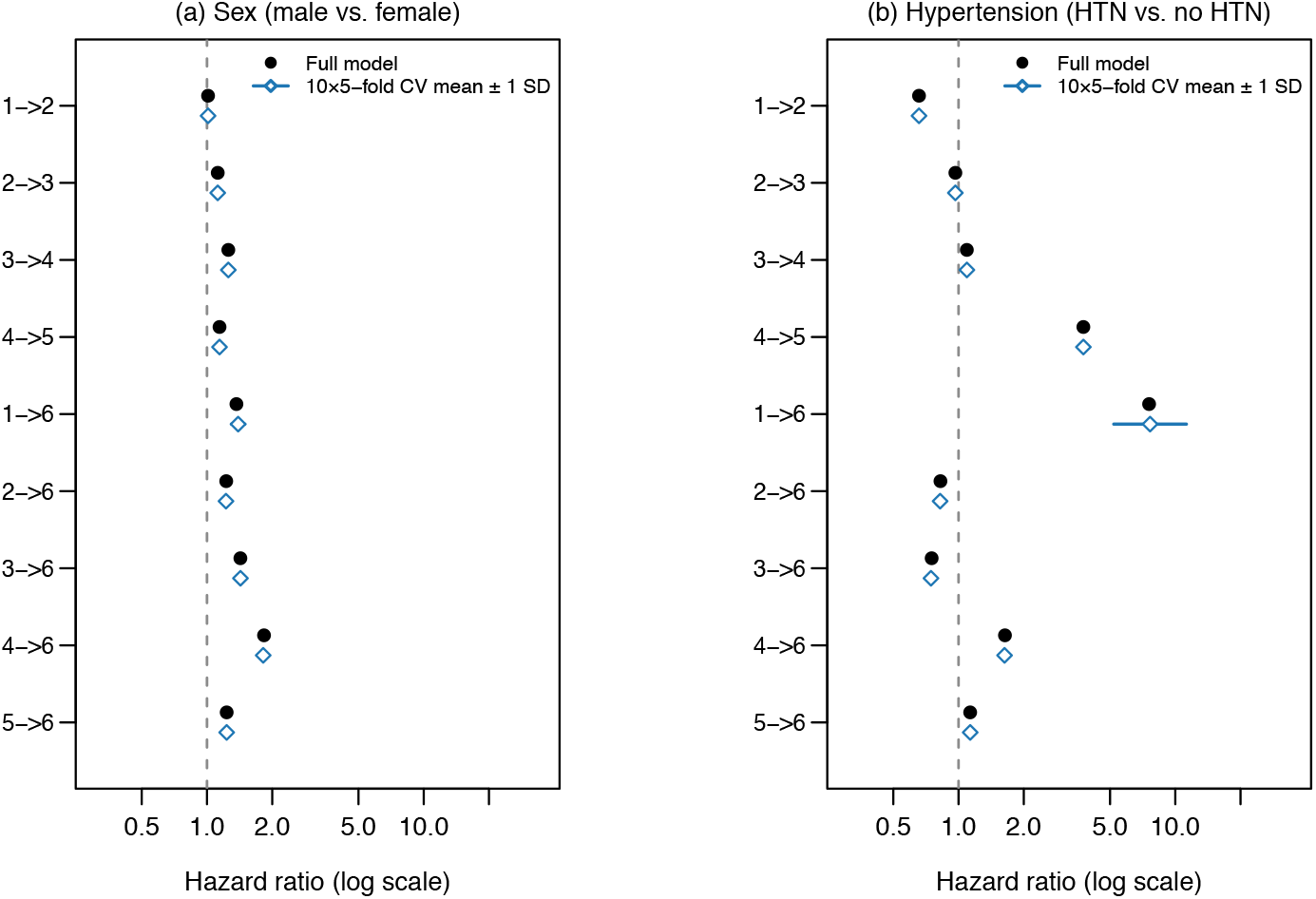
Forest plots of estimated hazard ratios for (a) sex (male vs. female) and (b) hypertension (HTN vs. no HTN). Filled circles show full-model point estimates; open diamonds with whiskers show cross-validation split-level (50 splits from 10×5-fold CV) means ±1 SD.

The CV split-level means (open diamonds) cluster tightly around the full-model estimates (filled circles) for almost all transitions. The notable exception is the 1 → 6 hypertension coefficient, which has a very wide confidence interval due to data sparsity and consequently shows greater variation across splits; this instability is examined further in the sensitivity analysis in Section 4.4. Male patients exhibit consistently faster disease progression and higher mortality than female patients across nearly all CKD stages in the simulated cohort. The only exception is the 1 → 2 transition, where sex has no statistically significant effect (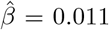, 95% CI: [ − 0.039, 0.062]). For intermediate disease progression, men move through Stages 2 → 3, 3 → 4, and 4 → 5 approximately 12% to 26% faster than women. In addition, men face significantly higher mortality hazards from every transient state, with hazard ratios ranging from 1.23 to 1.83. These findings show that sex has heterogeneous effects across transitions in a multi-state CTMC. The estimated hazard ratios are qualitatively consistent with patterns reported in epidemiological studies (Neugarten et al. 2000; Ricardo et al. 2019), suggesting that the fitted model produces transition patterns that are qualitatively consistent with prior epidemiological findings.

The estimated effect of hypertension on CKD progression varies by stage. In early stages, hypertension is associated with a slower 1 → 2 transition rate (HR = 0.66), while the 2 → 3 transition remains unaffected. At Stage 3, the effect becomes mixed: hypertension slightly increases the 3 → 4 transition rate (HR = 1.09) but is associated with lower mortality (HR = 0.75). The strongest effect appears at Stage 4, where hypertension substantially increases the transition to Stage 5 (HR = 3.77) and elevates mortality risk from Stages 4 and 5 (HR = 1.64 and 1.13, respectively). For the 1 → 6 transition, the estimated hazard ratio is large (HR = 7.57) but has a wide confidence interval [2.00, 28.62], reflecting data sparsity: among 212 observed Stage 1-to-Death transitions, only 4 occurred in patients without hypertension. One possible explanation is that the apparent protective effect in early stages reflects treatment-related factors, as suggested in prior clinical studies where hypertensive patients receive RAAS inhibitors that provide nephroprotection (Xie et al. 2016; Jafar et al. 2001). In later stages, this effect may be offset by cumulative structural damage associated with chronic hypertension (Klag et al. 1996; Fogo 2015). Although our analysis is based on synthetic data generated by a rule-based simulator and therefore cannot establish these mechanisms directly, this clinical interpretation provides one possible reading of the stage-specific pattern. It also illustrates how the multi-state CTMC framework can capture covariate effects that vary across disease stages.

### 4.4 Sensitivity analysis for the unstable parameter

As established in Section 4.3, the maximum likelihood estimator for the hypertension effect on the 1 → 6 transition is highly sensitive due to event sparsity. We conduct a formal sensitivity analysis to assess whether the model’s main findings depend on the precise value of this parameter.

We fit two constrained variants of the full model (*M* 1) by fixing *β*_htn,1→6_ at each boundary of its 95% confidence interval: *S*_lb_ fixes *β*_htn,1→6_ = 0.6945 (HR = 2.00, lower bound) and *S*_ub_ fixes *β*_htn,1→6_ = 3.3542 (HR = 28.62, upper bound). In both cases, the constrained coefficient is fixed, while all remaining estimable model parameters are re-estimated by maximum likelihood. Both constrained models converge successfully. Table 5 summarizes the parameter changes induced by fixing the unstable coefficient.

**Table 5.** Estimated coefficients 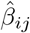 by fixing *β*_htn,1→6_ at its mean (Full) and lower (*S*_lb_) and upper (*S*_ub_) 95% CI bounds. Δ_max_ denotes the maximum relative change of the constrained estimate from the full-model estimate, defined as 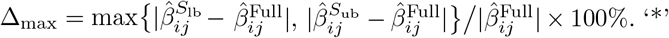 and ‘−’ symbols indicate coefficients are significant and not significant, respectively, for *α* = 0.05. The sole change in significance status is marked with †.

| Transition | Covariate | Full $\hat{\beta}_{ij}$ | $S_{lb}$ $\hat{\beta}_{ij}$ | $S_{ub}$ $\hat{\beta}_{ij}$ | $\Delta_{max}$ (%) | Full sig. | $S_{lb}$ sig. | $S_{ub}$ sig. |
| --- | --- | --- | --- | --- | --- | --- | --- | --- |
| 1 → 2 | sex | 0.0113 | 0.0113 | 0.0113 | 0.0% | — | — | — |
| 1 → 6 | sex | 0.3127 | 0.3074 | 0.3243 | 3.7% | * | * | * |
| 2 → 3 | sex | 0.1142 | 0.1142 | 0.1142 | 0.0% | * | * | * |
| 2 → 6 | sex | 0.2043 | 0.2063 | 0.2033 | 1.0% | * | * | * |
| 3 → 4 | sex | 0.2267 | 0.2267 | 0.2267 | 0.0% | * | * | * |
| 3 → 6 | sex | 0.3548 | 0.3543 | 0.3554 | 0.2% | * | * | * |
| 4 → 5 | sex | 0.1332 | 0.1332 | 0.1332 | 0.0% | * | * | * |
| 4 → 6 | sex | 0.6058 | 0.6010 | 0.6005 | 0.9% | * | * | * |
| 5 → 6 | sex | 0.2092 | 0.2094 | 0.2093 | 0.1% | * | * | * |
| 1 → 2 | htn | −0.4200 | −0.4191 | −0.4212 | 0.3% | * | * | * |
| 1 → 6 | htn | [fixed at CI bounds] | | | — | * | * | * |
| 2 → 3 | htn | −0.0336 | −0.0338 | −0.0338 | 0.6% | — | — | — |
| 2 → 6 | htn | −0.1924 | −0.1872 | −0.2061 | 7.1% | — | — | *† |
| 3 → 4 | htn | 0.0882 | 0.0884 | 0.0885 | 0.2% | * | * | * |
| 3 → 6 | htn | −0.2858 | −0.2869 | −0.2871 | 0.5% | * | * | * |
| 4 → 5 | htn | 1.3258 | 1.3260 | 1.3263 | 0.0% | * | * | * |
| 4 → 6 | htn | 0.4931 | 0.4986 | 0.4976 | 1.1% | * | * | * |
| 5 → 6 | htn | 0.1235 | 0.1241 | 0.1235 | 0.5% | * | * | * |

† Under $S_{ub}$ , the 2 → 6 htn estimate becomes marginally significant ( $\hat{\beta} = -0.206$ , 95% CI $[-0.407, -0.005]$ ).

Sex-related parameters remain stable. The maximum change across the nine sex coefficients is 3.7% (for the 1 → 6 transition), with all others changing by at most 1%. All significance conclusions for sex are preserved under both constrained scenarios. Other hypertension parameters are robust. Among the eight freely estimated hypertension coefficients, the maximum change is 7.1% (for the 2 → 6 transition). This specific coefficient (−0.1924 in the full model) crosses the significance threshold only under the extreme upper-bound scenario *S*_ub_, where it shifts to − 0.2061 and becomes marginally significant. All other hypertension significance conclusions—including the 4 → 5 acceleration (HR = 3.77), the apparent 1 → 2 protective association, and the apparent 3 → 6 protective association—are fully robust to the constraint. Overall, this sensitivity check suggests that the main interpretive findings of the fitted model do not depend strongly on the precise value of the sparse 1 → 6 hypertension coefficient.

### 4.5 Clinical interpretation and application

While the estimated hazard ratios (Sections 4.3 and 4.4) quantify instantaneous relative risks, interpreting the macroscopic behavior of the multi-state CTMC model requires evaluating absolute time. To translate these transition rates into tangible cohort projections, Table 6 reports the expected mean sojourn times (years spent in each stage).

**Table 6.** Estimated mean sojourn times (years spent in each stage before transitioning) by covariate profile. Values are point estimates with 95% confidence intervals in brackets.

| Profile | Stage 1 | Stage 2 | Stage 3 | Stage 4 | Stage 5 |
| --- | --- | --- | --- | --- | --- |
| Female, no HTN | 1.84 [1.71, 1.98] | 5.51 [5.20, 5.84] | 4.69 [4.42, 4.97] | 3.28 [3.08, 3.49] | 13.40 [12.22, 14.70] |
| Male, no HTN | 1.82 [1.70, 1.95] | 4.88 [4.61, 5.16] | 3.69 [3.48, 3.91] | 2.78 [2.62, 2.95] | 10.87 [9.97, 11.86] |
| Female, HTN | 2.72 [2.62, 2.83] | 5.77 [5.58, 5.97] | 4.43 [4.28, 4.58] | 0.90 [0.87, 0.93] | 11.85 [11.26, 12.46] |
| Male, HTN | 2.67 [2.58, 2.76] | 5.11 [4.98, 5.25] | 3.49 [3.41, 3.58] | 0.78 [0.76, 0.80] | 9.61 [9.29, 9.94] |

The hazard ratios from Section 4.3 directly affect how many years a patient spends in each CKD stage. Without hypertension, simulated men spend approximately 0.5 to 1.0 fewer years in each intermediate stage (Stages 2, 3, and 4) than women. Hypertension shifts these timelines even more substantially: for female profiles, it adds nearly a full year to Stage 1, while cutting the time spent in Stage 4 by more than two-thirds, from 3.28 years to 0.90 years. Furthermore, we plot the total expected survival and long-term survival probabilities in Figure 5.

**Fig. 5.**
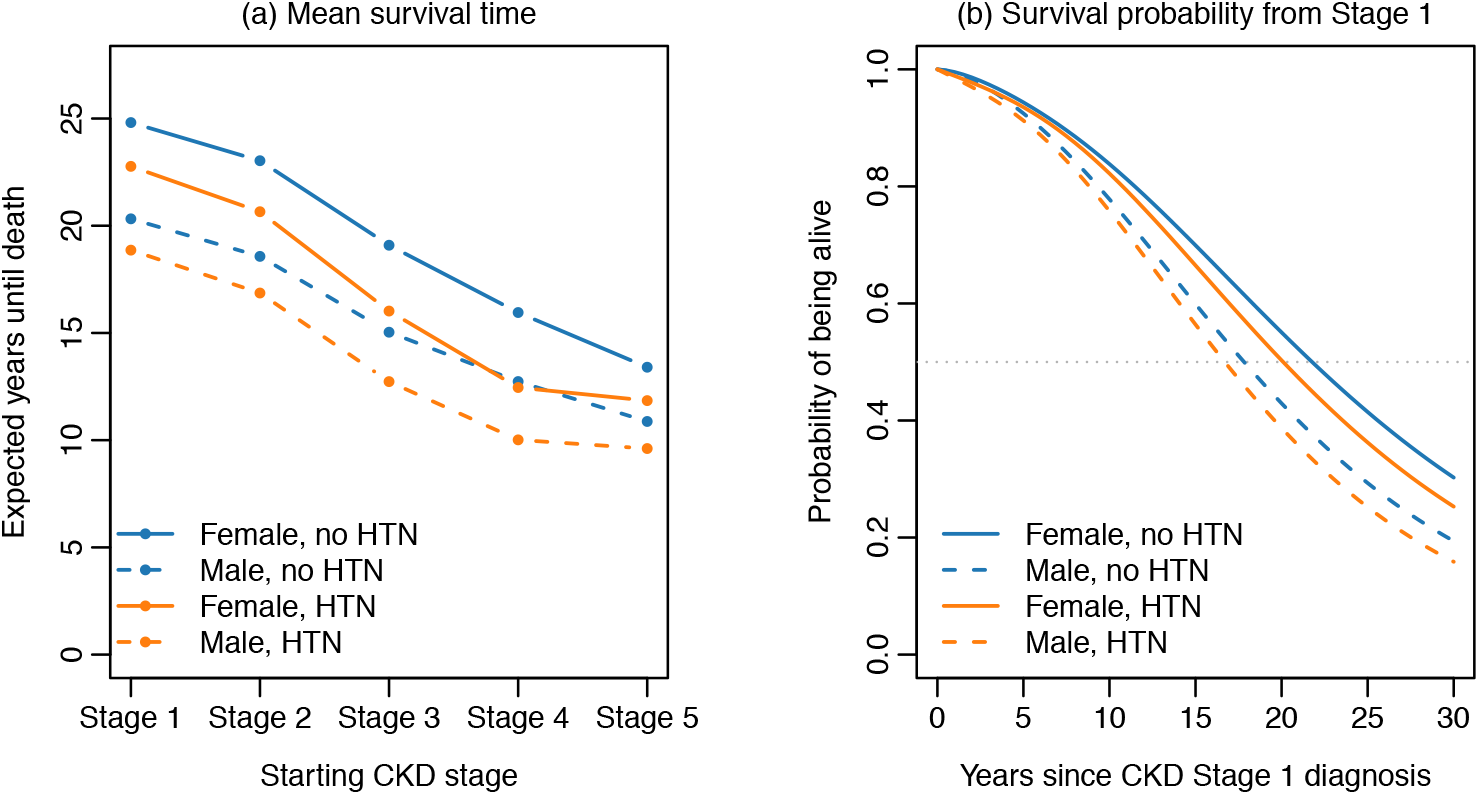
(a) Expected years until death by starting CKD stage and covariate profile. (b) 30-year survival probability curves for a patient diagnosed at CKD Stage 1, by covariate profile.

Figure 5a shows that sex is a major driver of long-term survival in the synthetic cohort, with female profiles having consistently longer expected survival than male profiles across all starting CKD stages. Figure 5b shows the same pattern over a 30-year horizon: for patients diagnosed at Stage 1, females reach median survival at roughly 21 to 22 years, whereas males reach it at approximately 16 to 17 years. Hypertension also has a negative net effect on projected survival. In Figure 5a, hypertensive patients have consistently shorter expected survival than non-hypertensive patients within the same sex group across early and intermediate stages. Thus, although hypertension prolongs expected sojourn time in some earlier stages within the model, this does not offset the accelerated decline at later stages. This study is based on synthetic data generated by a rule-based simulator. While such data provide a controlled setting for illustrating the proposed modeling framework, the observed patterns may reflect the underlying data-generating assumptions rather than real-world clinical mechanisms. Therefore, the covariate effects discussed here should be interpreted as model-based examples rather than clinical conclusions.

## 5 Conclusion

This work presented a unified six-state CTMC framework for modeling CKD progression, with CKD stages 1–5 represented as transient states and death as an absorbing state. We considered both the general non-homogeneous setting, where transition rates may vary with time, and the time-homogeneous setting, where the transition rate matrix is constant. Within this framework, we derived transition probabilities, state distributions, sojourn times, survival-related quantities, and expected patient counts, making explicit a set of quantities that are often used implicitly in applied multi-state analyses.

In the empirical analysis, we focused on the homogeneous model as a tractable baseline and fitted it to synthetic longitudinal EHR data. The results showed that the covariate model improved predictive fit relative to the null model, that the fitted CTMC captured the main population-level prevalence patterns, and that covariate effects were transition-specific rather than uniform across CKD stages. These findings illustrate how a multi-state CTMC framework can separate stage-to-stage progression, mortality, and covariate effects within a single disease progression model. Because the analysis is based on synthetic data, these patterns should be interpreted as a modelbased assessment of the framework rather than as clinical evidence about real-world CKD progression.

The derived sojourn-time and survival summaries further demonstrate how transition-rate estimates can be translated into interpretable descriptions of CKD trajectories. In this way, the framework links local transition behavior with longer-term quantities such as stage occupancy, time spent in each disease stage, and expected survival time.

Several extensions remain of interest. A fully non-homogeneous implementation may better capture age-dependent progression patterns, while hidden multi-state formulations may help account for measurement noise and short-term fluctuations in observed eGFR-based disease stages. More broadly, the framework provides a mathematically explicit starting point for future CTMC-based CKD progression studies involving both stage transitions and survival outcomes, especially when applied to real longitudinal clinical data.

## Declarations

### Ethics Approval

Not applicable; only synthetic data were used.

### Consent to Participate

Not applicable.

### Consent for Publication

Not applicable.

### Data Availability

The synthetic dataset and processed panel are available at https://github.com/ShahriyariLab/Multi-state-Continuous-Time-Markov-Chain-Modeling-for-Chronic-Kidney-Disease-Progression; generation parameters are given in Appendix C.

### Code Availability

All analysis code is available at https://github.com/ShahriyariLab/Multi-state-Continuous-Time-Markov-Chain-Modeling-for-Chronic-Kidney-Disease-Progression.

### Author Contributions

All authors contributed to the conception, analysis, and writing of this manuscript.

## Appendix A Transition Probability Functions

In this appendix, we provide explicit closed-form expressions for the transition probability functions *P*_*ij*_(*τ, t*) under the homogeneous CTMC assumption. We first write down the forward Kolmogorov ODE system induced by the six-state structure and then solve it under the assumption that the relevant exit rates are distinct. To illustrate how repeated exit rates can be handled, we work out the case of *P*_12_(*τ, t*) when *λ*_1_ = *λ*_2_ and verify that the same expression is obtained by taking the limit of the distinct-rate formula. Other repeated-rate cases can be treated similarly, although the explicit forms depend on the particular multiplicity pattern. These formulas support the transition probability expressions used in the main text under the homogeneous baseline model.

For the homogeneous CTMC, from the Kolmogorov forward equations, we have

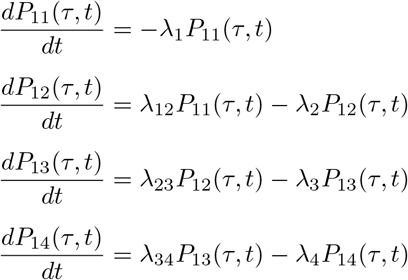

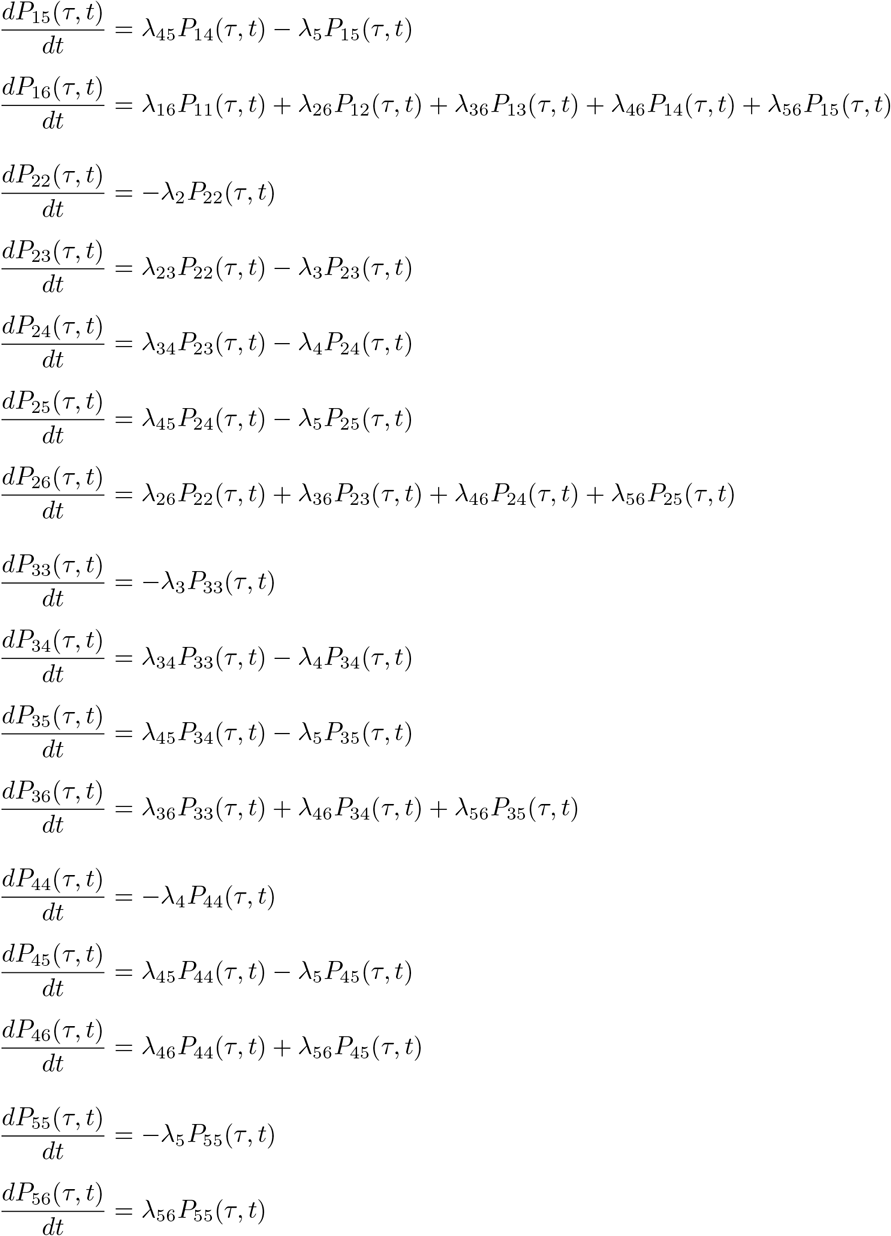

where *λ*_*i*_ = ∑_*j* ≠ *i*_ *λ*_*ij*_ represents the total exit rate from state *i*, specifically *λ*_1_ = *λ*_12_ + *λ*_16_, *λ*_2_ = *λ*_23_ + *λ*_26_, *λ*_3_ = *λ*_34_ + *λ*_36_, *λ*_4_ = *λ*_45_ + *λ*_46_, and *λ*_5_ = *λ*_56_.

We can calculate an analytic expression for each element of *P* (*τ, t*) by solving these equations. When the exit rates *λ*_*i*_, …, *λ*_*j*_ are distinct, we yield the following explicitm forms

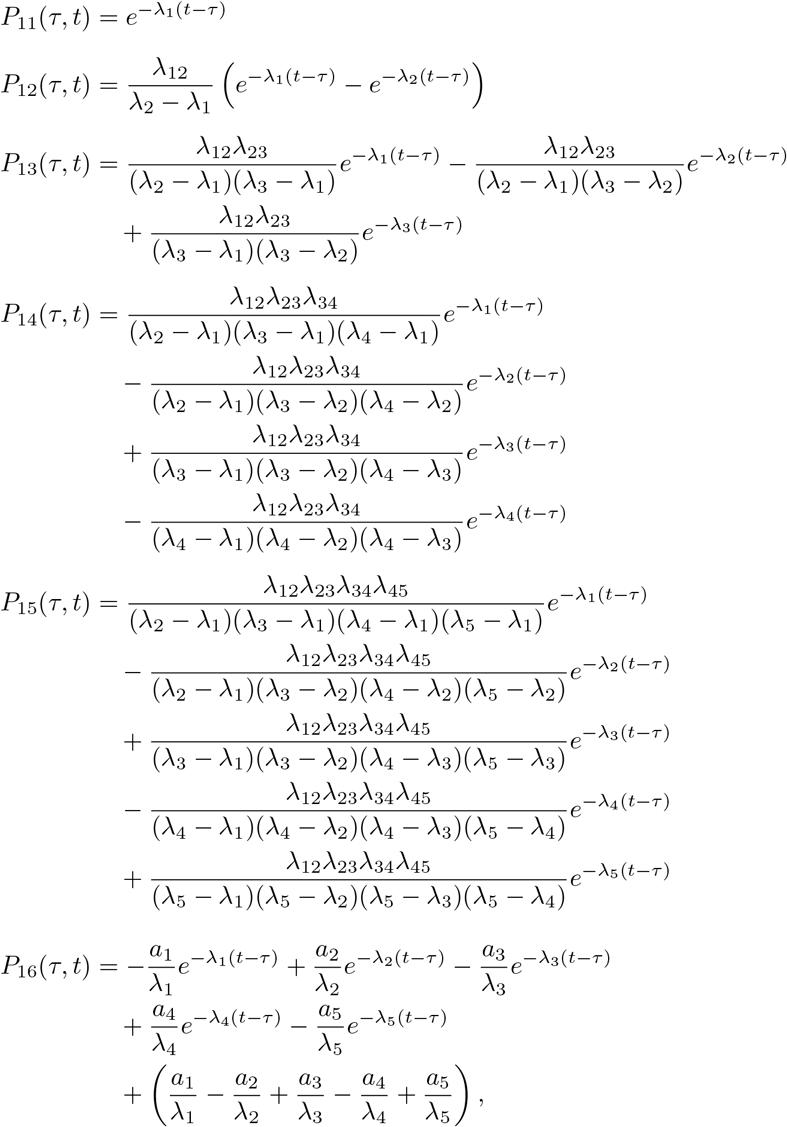

where the coefficients *a*_*i*_ are defined as

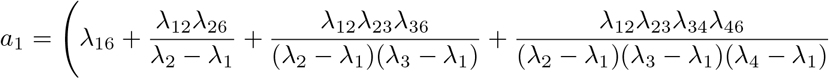

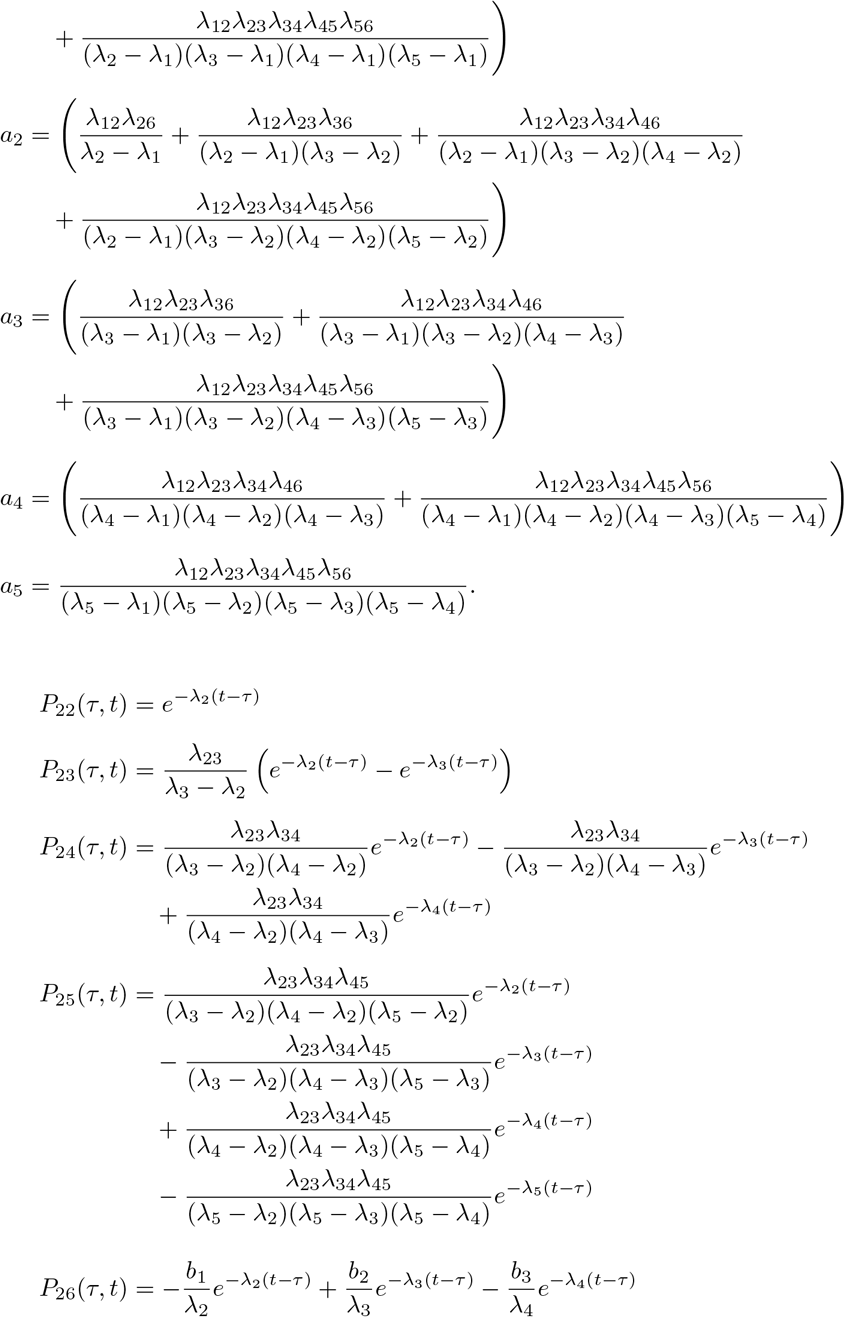

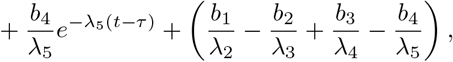

where the coefficients *b*_*i*_ are defined as

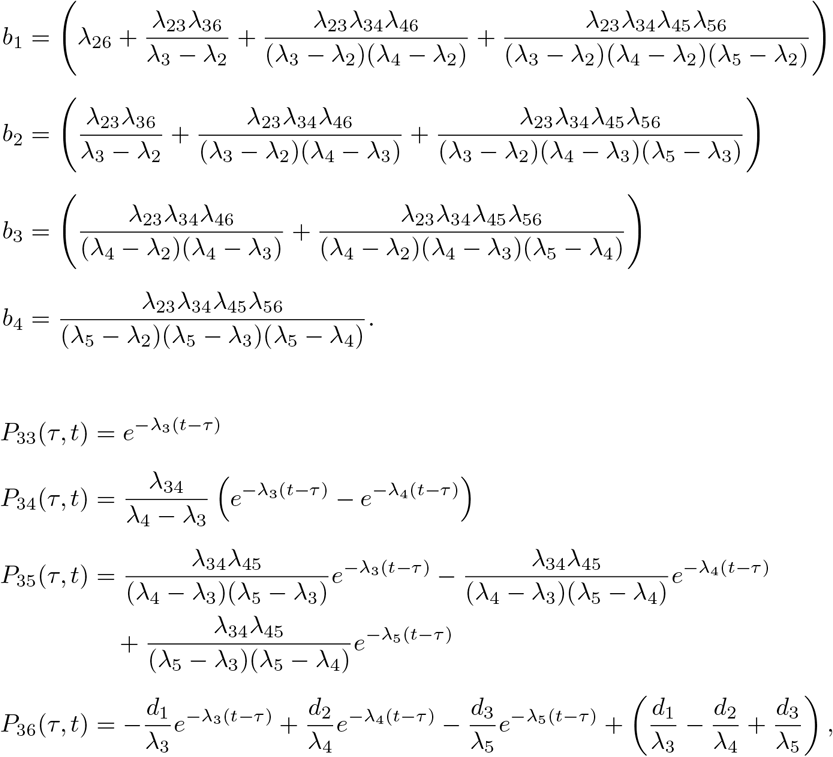

where the coefficients *d*_*i*_ are defined as

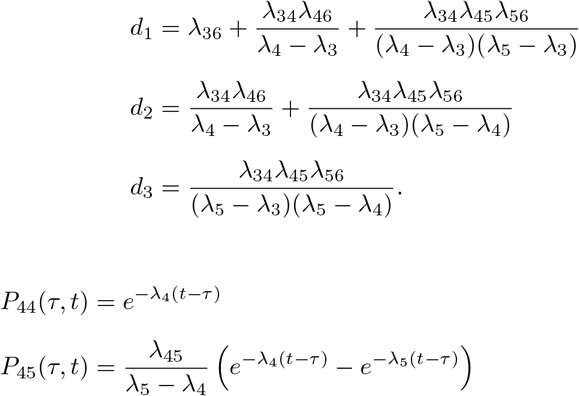

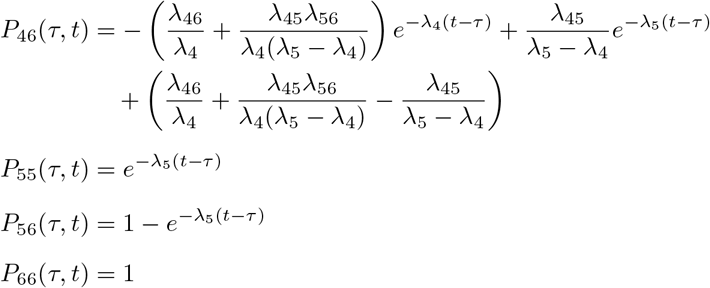

When two exit rates coincide, the compact partial-fraction expression should be replaced by the corresponding repeated-root solution. For example, consider *P*_12_(*τ, t*). Since 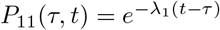, the forward equation for *P*_12_ is

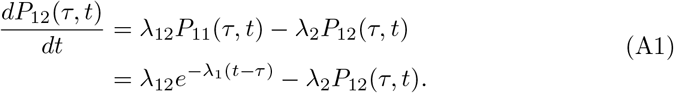

If *λ*_1_ = *λ*_2_ = *λ*, then (A1) becomes

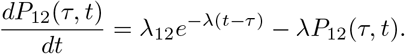

Multiplying both sides by the integrating factor *e*^*λt*^, we obtain

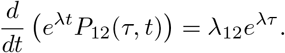

Integrating from *τ* to *t* and using *P*_12_(*τ, τ* ) = 0 in (8), we have

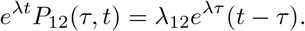

Therefore,

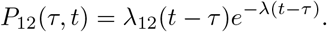

The same expression can also be obtained as the limiting value of the distinct-rate formula (10). When *λ*_1_≠ *λ*_2_,

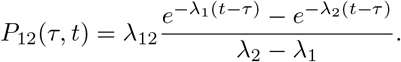

Taking the limit as *λ*_2_ → *λ*_1_ = *λ*, we obtain

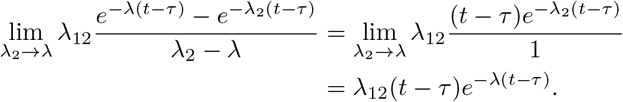

Thus, the repeated-rate solution agrees with the continuous limit of the distinct-rate expression.

## Appendix B State Probability Distribution

In this appendix, we give the explicit analytical expressions of the state probability distributions *p*_*j*_(*t*) for the homogeneous CTMC. Following the same pattern as Appendix A, we start from the population-level forward Kolmogorov ODEs and present the closed-form solution under distinct exit rates. The repeated-rate case is illustrated using *p*_2_(*t*) as a worked example; the remaining repeated-rate expressions follow by the same limiting argument. These formulas describe how the proportion of patients in each CKD stage evolves over time and underpin the prevalence and expected-count results reported in the main text.

For the homogeneous CTMC, from the Kolmogorov forward equations, we have

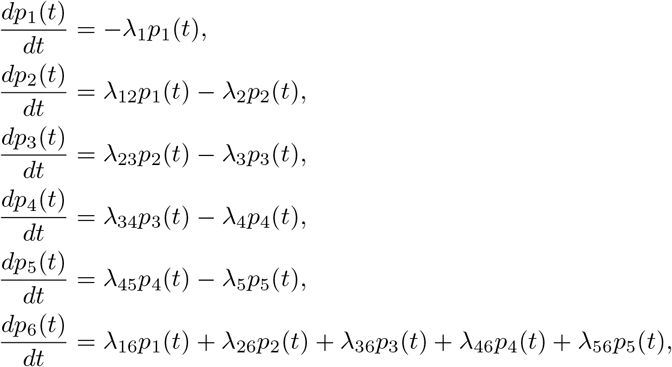

where *λ*_1_ = *λ*_12_ + *λ*_16_, *λ*_2_ = *λ*_23_ + *λ*_26_, *λ*_3_ = *λ*_34_ + *λ*_36_, *λ*_4_ = *λ*_45_ + *λ*_46_, and *λ*_5_ = *λ*_56_.

By solving these differential equations, when the exit rates *λ*_*i*_, …, *λ*_*j*_ are distinct, we obtain the exact analytical expressions

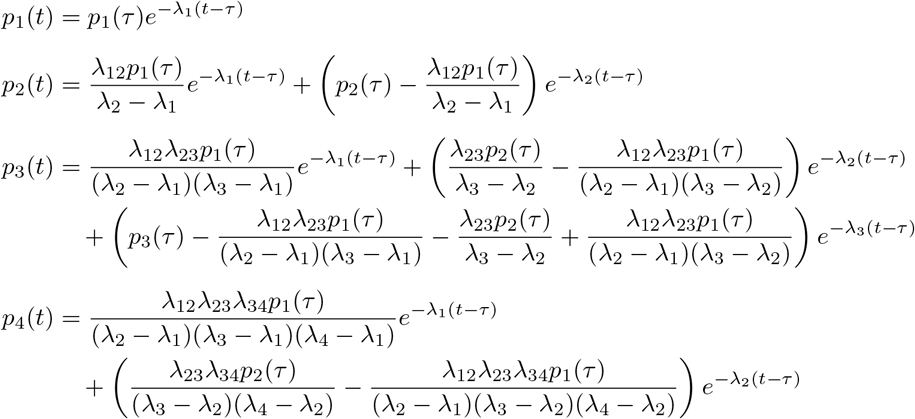

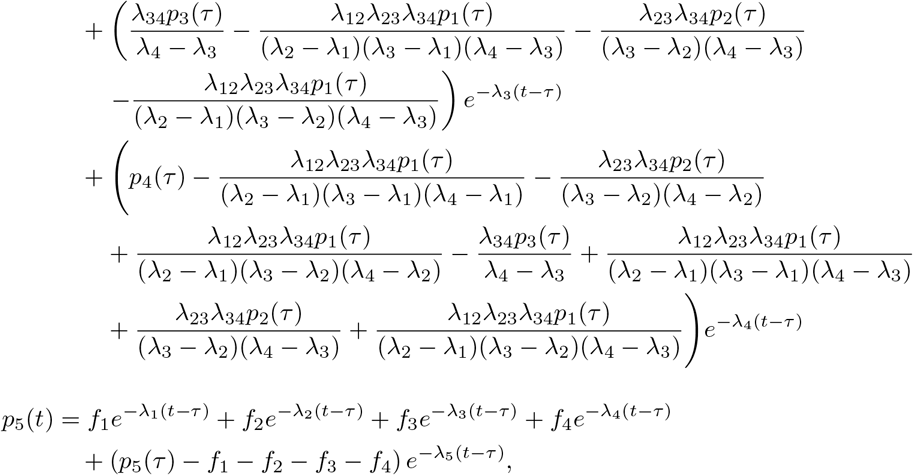

where the coefficients *f*_*i*_ are defined as

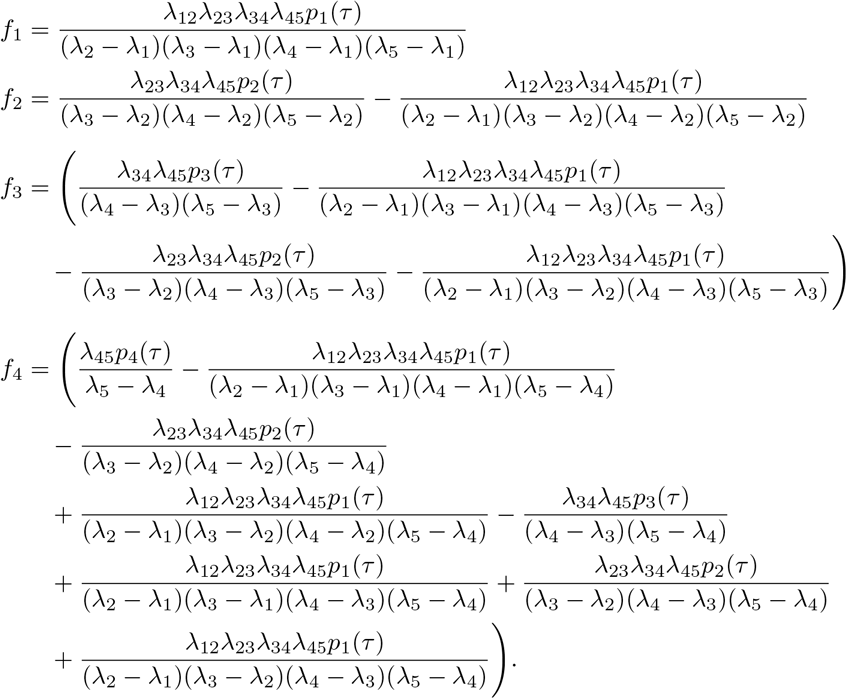

Finally, for the absorbing state, we have

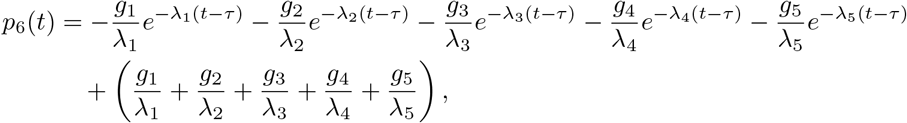

where the coefficients *g*_*i*_ are defined as

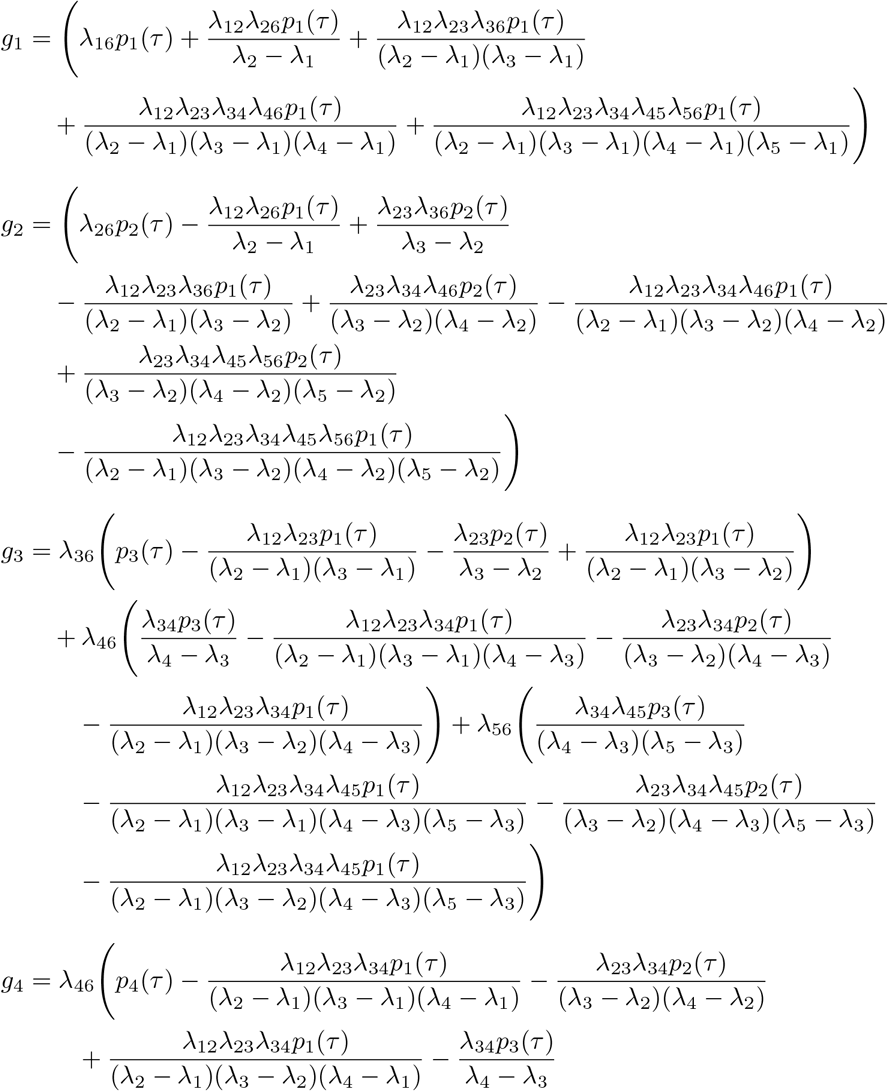

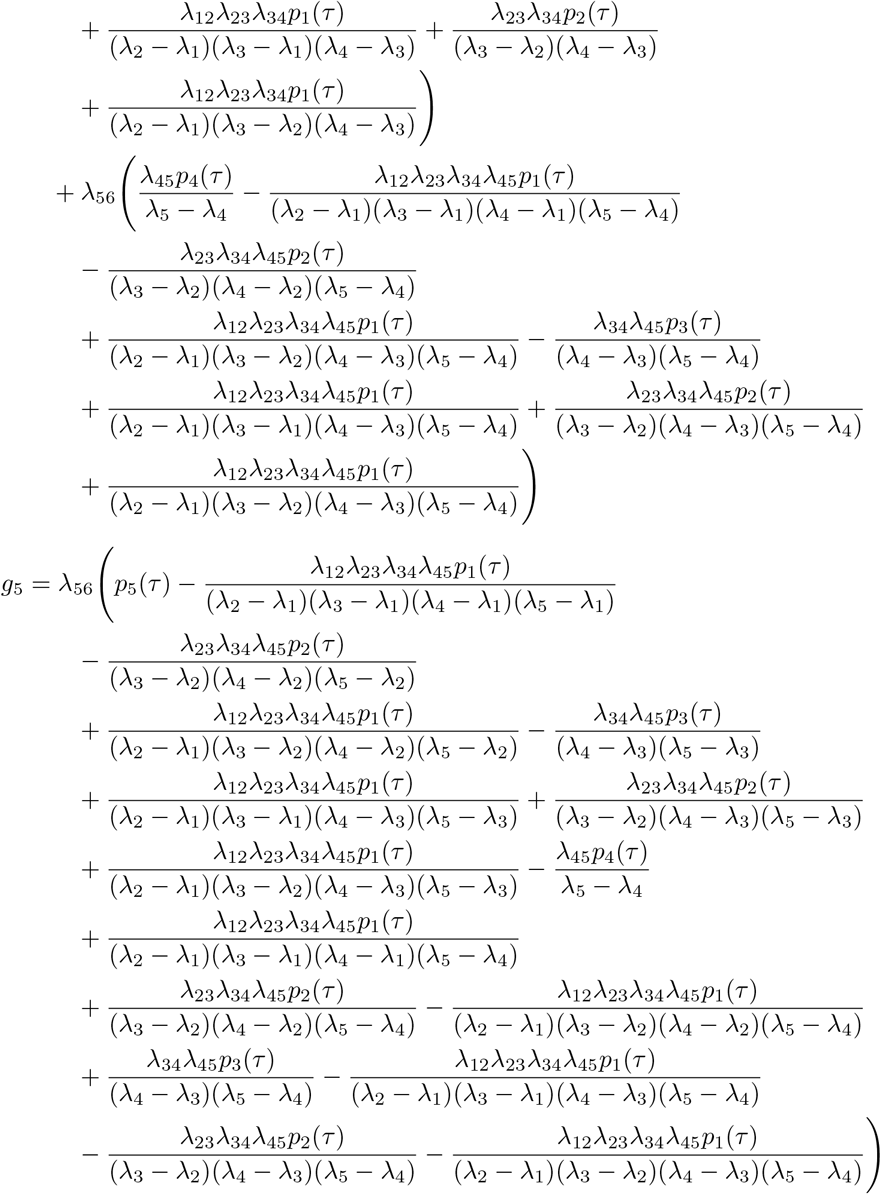

When two exit rates coincide, the compact partial-fraction expression should be replaced by the corresponding repeated-root solution. For example, consider the equation for *p*_2_(*t*):

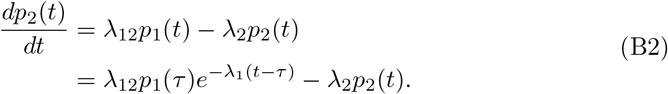

When *λ*_1_ = *λ*_2_ = *λ*, the equation (B2) becomes

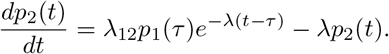

Multiplying both sides by the integrating factor *e*^*λt*^, we obtain

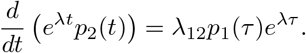

Integrating from *τ* to *t* gives

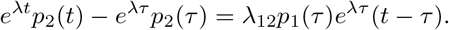

Therefore,

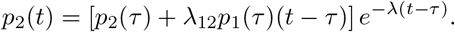

This repeated-rate solution can also be obtained by taking the limit of the distinct-rate expression as *λ*_2_ → *λ*_1_ = *λ*. Indeed,

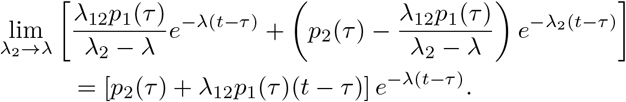

## Appendix C Implementation details for the empirical analysis

This appendix documents the data generation, preprocessing, model fitting, cross-validation, and prevalence calculation procedures used in Sections 4.1 and 4.2.

### C.1 Synthetic data generation

The synthetic cohort is produced with Synthea v4.0.0 (Walonoski et al. 2018) using random seed 1219 and a target population of 200,000. Synthea writes its longitudinal output as a set of CSV files following a fixed EHR schema, including `patients.csv` (demographics, birth and death dates), `conditions.csv` (diagnoses with onset dates), and `observations.csv` (lab values and vital signs with LOINC codes). We retain the 21,100 patients (9.2%) of the 229,989 simulated individuals who receive at least one Chronic Kidney Disease diagnosis in the Synthea-generated `conditions.csv`. eGFR measurements are read from the Synthea-generated `observations.csv` using LOINC codes 33914-3 (MDRD) and 62238-1 (CKD-EPI).

### C.2 Cohort construction and preprocessing

The longitudinal panel used for model fitting (`ckd_panel.csv`, 335,283 rows from 21,100 patients) is built from the Synthea CSV output by our preprocessing script `preprocess.py` in the following steps.

1. **eGFR-to-state mapping**. Each eGFR value is mapped to a CKD stage 1–5 by the KDIGO thresholds in Table 1.
2. **Per-day deduplication**. If a patient has multiple eGFR measurements on the same date, keep the one corresponding to the most severe stage.
3. **Death rows**. For each deceased patient, append a single observation at the date of death with state = 6.
4. **Monotonicity**. For each patient, replace the state sequence by its running maximum (cumulative max). This enforces the irreversibility assumption of the CTMC; in practice, downward fluctuations in eGFR-derived stage are treated as measurement noise around the true progression.
5. **Six-month thinning with transition preservation**. Within each six-month age bin, if the patient’s state does not change, only the first record is kept; if a transition occurs in the bin, the last record before the transition and the first record after are both retained. All observed transitions are therefore preserved.
6. **Minimum two observations**. Patients with only one record (no observable transition) are excluded.

### C.3 Cross-validation procedure

The 10-times-repeated 5-fold cross-validation produces 50 train/test splits. The split is performed at the patient level, stratified on the initial CKD stage, and each of the 10 repeats uses a fresh random partition seeded by 42 + *r* for *r* = 1, …, 10. For each (repeat, fold) pair we fit the full model *M* 1 and the null model *M* 0 on the 80% training set and evaluate the held-out *LL/n* on the remaining 20%. A small number of folds occasionally fail to converge from the default data-estimated starting values; in those cases we refit using alternative starting values until convergence is achieved (3 of 50 folds in our run required this). All implementation details—optimisation parameters, the convergence-recovery strategy, the parallel-execution setup, and the significance test on held-out *LL/n*—are available in the public repository (see Appendix C.5).

### C.4 Prevalence calculation algorithm

Both the empirical and the model-predicted prevalences in Section 4.2 are computed by the same routine, `compute_prev()`. Fix a window half-width *w* = 2.5 years. For each target age *t* ∈ {40, 45, …, 85} and each patient *k*:

1. **Anchor observation**. Let *τ*_*k*_ be the age of the most recent record of patient *k* prior to *t* − *w*, and let *s*_*k*_ be the observed state at *τ*_*k*_. If no such record exists, *k* is skipped (no baseline available).
2. **Lost-to-follow-up check**. If *k* has not been observed to have died at any age ≤ *t* and *k*’s last record occurs strictly before *t* − *w*, then *k* is skipped (lost to follow-up).
3. **Observed state** *ŷ*_*k*_(*t*). For the patients *k* that pass steps 1–2, *ŷ*_*k*_(*t*) is assigned by the following priority:
  - if *k* has died at any age ≤ *t*, set *ŷ*_*k*_(*t*) = 6;
  - else if *k* has at least one observation in [*t* − *w, t* + *w*], set *ŷ*_*k*_(*t*) equal to the state of the observation closest to *t*;
  - else, set *ŷ*_*k*_(*t*) = *s*_*k*_ (last-observation-carried-forward).
4. **Predicted distribution**. If *s*_*k*_ = 6, the predicted distribution is the point mass on state 6. Otherwise, the predicted distribution is the row vector 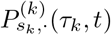 of transition probabilities from *s*_*k*_, evaluated at the fitted model with patient *k*’s covariates.

#### Remark

The strict requirement *τ*_*k*_ *< t* − *w* in step 1, combined with the lost-to-followup check in step 2, ensures that the prediction horizon *dt* = *t* − *τ*_*k*_ is at least *w* years. This guarantees that the model produces a genuine forward prediction rather than predicting the state from a near-contemporaneous observation.

Let 𝒦_*t*_ denote the set of patients retained after steps 1–2. The empirical and model-predicted prevalences at age *t* are

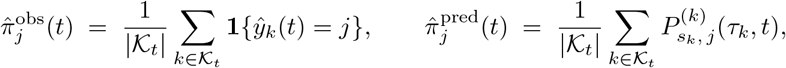

for *j* = 1, …, 6. The two share the same denominator |𝒦_*t*_| and are therefore directly comparable. The full-data prevalence curves in Figure 3 (solid lines) use 𝒦_*t*_ defined on the full cohort; the OOS curves (dashed lines) compute 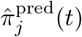 within each held-out fold using only that fold’s evaluable patients and then average across the 50 splits, with ±1 SD bands across splits.

### C.5 Code availability and software

All preprocessing, model fitting, cross-validation, and prevalence-calculation code is available at https://github.com/ShahriyariLab/Multi-state-Continuous-Time-Markov-Chain-Modeling-for-Chronic-Kidney-Disease-Progression. R analyses use R 4.4 with the `msm` (Jackson 2011) and `parallel` packages; preprocessing uses Python 3 with `pandas` and `numpy`. Random seeds for the Synthea generation and the cross-validation partitions are fixed in the code to ensure reproducibility.

